# Non-mulberry silk fibroin functionalization enhances charge-transfer efficiency in aligned polypyrrole-silk composites for electrically stimulated neurite outgrowth

**DOI:** 10.1101/2024.12.12.628243

**Authors:** Rajiv Borah, Joseph Christakiran Moses, Jnanendra Upadhyay, Jitu Moni Das, Biman B. Mandal

## Abstract

Electroconductive biomaterials (ECBs) replicate the natural bioelectrical environment of nerve tissue, promoting action potential propagation after injury and enhancing nerve regeneration through therapeutic electrical stimulation (ES). We present a highly electroactive Faradaic ECB with exceptional electrical conductivity and charge density, alongside low electrochemical impedance. These ECBs trigger action potentials at low stimulation voltages by regulating redox reactions through their intrinsic reversible behavior, thereby preventing electrode degradation and tissue damage. Our biohybrid scaffold consists of aligned microfibrous matrices of polypyrrole (PPy) and *Bombyx mori* silk fibroin (BmSF), functionalized with *Antheraea assamensis* silk fibroin (AaSF) rich in the cell-affinitive RGD tripeptide. Serving as an anionic dopant for PPy, AaSF significantly enhances the scaffold’s electrical properties (∼9.18 mS cm^-1^) and charge-transfer efficiency (∼25.27 Ω). The scaffolds exhibit superior charge injection capacity at low potentials compared to conventional bioelectrodes (e.g., 0.46 mC cm^-2^ at 50 mV). Under pulsed ES at 50 mV cm^-1^, these scaffolds support remarkable neurite outgrowth of dorsal root ganglion (DRG) neurons up to 830 μm (7 days). Notably, higher current densities and voltages decrease the rate of neurite outgrowth, highlighting the importance of optimizing ES parameters to effectively evoke functional action potentials without causing any neuronal damage. Biocompatibility assessments reveal that AaSF functionalization improves cellular behavior while minimizing immunomodulatory responses. Enhanced neuronal and glial differentiation is attributed to better cell communication facilitated by excellent adhesion and increased conductivity. In essence, this study provides a strategy for selecting optimal ES parameters for electrically excitable tissues using established electrochemical techniques. The fabricated biohybrid scaffolds hold significant promise as smart nerve guidance channels (NGCs) for future nerve regeneration therapies.

## 1. Introduction

The peripheral nervous system (PNS), consisting of sensory and motor neurons, is an electrosensitive tissue having conductivity ranging between 0.00008 Scm^-1^ to 0.001 S cm^-1^.^1^ Therefore, biomaterials intended for nerve repair should ideally be electroconductive to mimic the native bioelectric microenvironment. Moreover, the application of external electrical stimulation (ES) through such electroconductive biomaterials (ECBs) significantly enhances nerve regeneration processes^2, 3^. Any uncontrolled change in membrane polarization during ES through ECBs can result in dysfunction in voltage-gated channels of neurons leading to neuronal excitotoxicity and death. This highlights the importance and the need for an effective and safe ES protocol, which can be achieved by stimulating cells and tissues at low potentials using ECBs with high charge density and charge-transfer efficiency^4, 5^.

Previous studies indicate that ECBs should inherently exhibit enhanced electroactivity for efficient charge transfer, allowing membrane depolarization at low stimulation voltages within safe limits for living tissues.^5–7^ To achieve this, ECBs must have adequate conductivity, charge storage capacity (CSC), charge/current injection capacity, and low electrochemical charge transfer resistance (R_ct_). The CSC, which is the available charge density within a voltage range, should be high to ensure adequate charge is available for membrane depolarization during ES.^5, 7^ Additionally, the charge-transfer reactions at the ECB and tissue interface should facilitate the transition from electron flow to ion flow in the tissue, necessitating a low R_ct_ to make the available charges in ECBs easily accessible. ECBs support charge/current injection through capacitive and faradaic mechanisms. Conventional capacitive neural electrode materials (e.g., titanium nitride (TiN), tantalum/tantalum oxide, etc.) store and transfer charge through the charging and discharging of the electrode-electrolyte double layer.^7^ While these reactions are stable and do not form chemical species during ES, they result in lower charge density, limiting the ability to achieve high charge-injection capacity. Conversely, Faradaic materials like electrically conducting polymers (ECPs) demonstrate significant charge storage capacity through their intrinsic oxidation and reduction reactions at the electrode-electrolyte interface for charge transfer. ECPs facilitate both electron and ion transport during ES, offering a high level of charge for stimulation. However, there is a risk of irreversible redox reactions leading to electrode degradation and tissue damage, which can be mitigated by selecting low stimulation potential and utilizing charge injection waveforms with appropriate parameters. Therefore, we propose that a highly electroactive Faradaic ECB, characterized by high electrical conductivity, charge density, and low electrochemical impedance, may trigger an action potential for membrane depolarization at low stimulation potentials.

Polypyrrole (PPy) stands out as one of the most extensively researched ECPs for tissue engineering applications.^3, 8, 9^ PPy demonstrates exceptional electrochemical and conductive properties, alongside *in vitro* and *in vivo* biocompatibility.^2, 8, 10^ However, creating aligned scaffolding matrices mimicking the neural anatomy to guide axonal regeneration using PPy is challenging due to its limited solubility. Nevertheless, PPy offers the advantage of straightforward conjugation with biologically active molecules^9^. To address this, we chose *Bombyx mori* silk fibroin (BmSF) fibers as a scaffolding matrix to compliment the PPy’s conductive properties. [**Figure 1**]. BmSF has been documented to be biocompatible, minimally immunogenic, confers tunable biodegradability and bestows chemical reactivity for further functionalization by virtue of the presence of reactive amino acids (such as lysine, tyrosine, serine, glutamate etc.).^11, 12^

**Figure 1:**
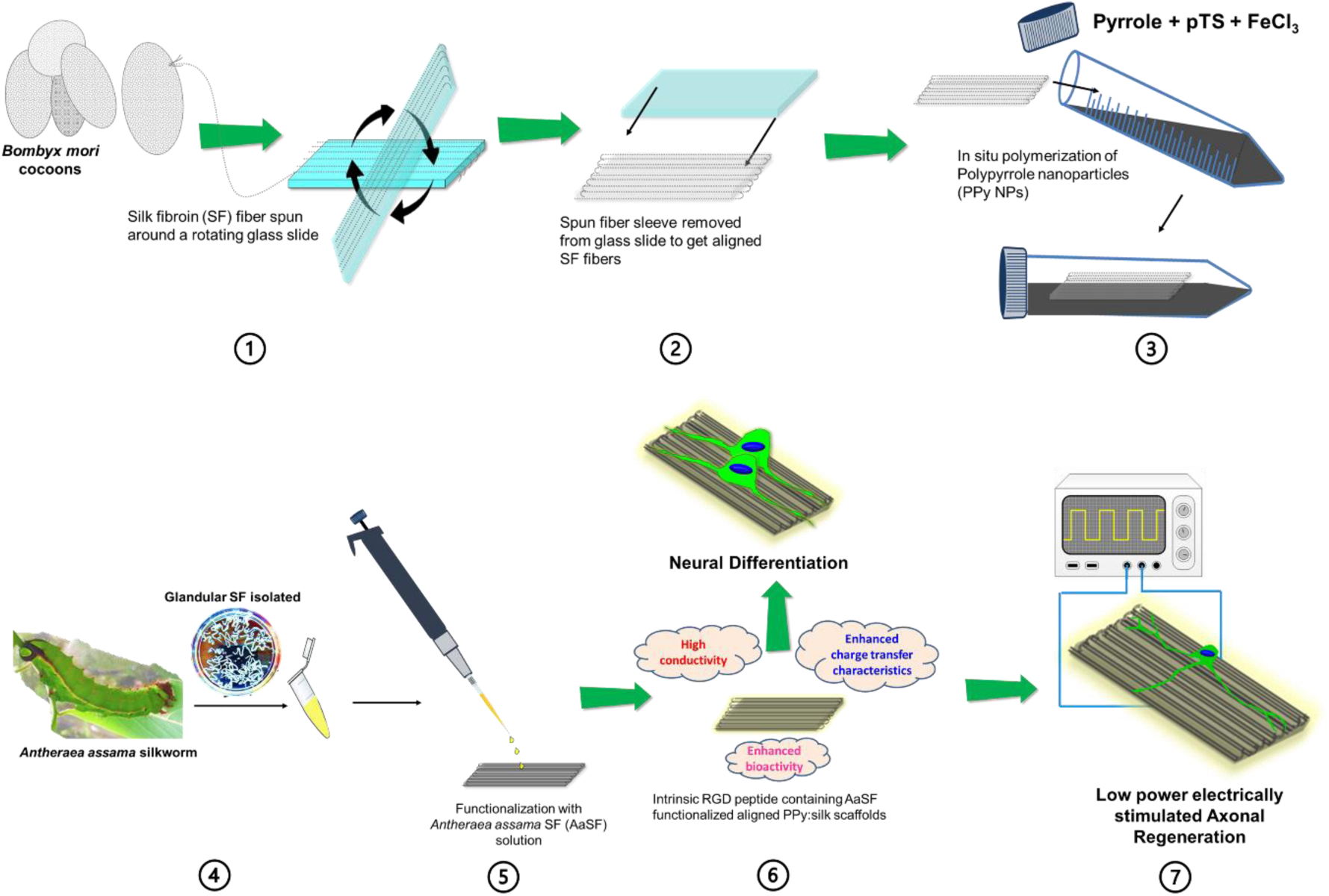
Scheme illustrating the fabrication of highly conductive PPy:Silk based aligned scaffolds with higher charge storage capacity and lower charge transfer resistance and its efficacy evaluation in terms of electrically stimulated neurite outgrowth at low stimulation potential.

Previous studies have demonstrated that conductive PPy-incorporated silk-based biomaterial scaffolds can be developed in various material formats, including hydrogels, sponges, films, fibers, and 3D-printed conduits.^8, 10, 13–19^ **Table S1** (Supporting information) lists the salient features of these studies which apply different fabrication approaches for obtaining aligned fibers including electrospinning and 3D printing. In most of the studies, PPy was incorporated onto silk scaffolds as a coating through chemical polymerization.^8, 10, 16–19^ Existing research emphasizes the need for a detailed investigation of the electrochemical behavior of PPy:Silk scaffolds to uncover their potential link to electrically stimulated neurite outgrowth. Our approach here was to utilise a very facile fabrication method to obtain aligned silk fibers (Figure 1). For this purpose, we chose to use BmSF fibers as the backbone and on which PPy was coated to confer electroconductive traits. Furthermore, to improve the bioactivity of these silk-PPy matrices we functionalized it with SF protein extracted from the Indian non-mulberry *Antheraea assamensis* (Aa) silkworm, which contains the cell-affinitive RGD tripeptide (Arg-Gly-Asp). The inclusion of (AaSF) also is envisaged to improve the mechanical properties due to its poly(alanine) sequences.^11^ The superior biological performance of AaSF based biomaterials have been already demonstrated in cardiac, bone, skin, and cartilage tissue regeneration both *in vitro* and *in vivo*.^20^ This study, for the first time, investigates the potential of AaSF in nerve growth. It also aims to assess the impact of the anionic nature of AaSFon the electroactivity of PPy.

We comprehensively investigated the effect of AaSF functionalization on the physiochemical properties of the PPy:Silk scaffold, such as electroconductivity, mechanical properties, and surface shape. To assess the performance of the developed matrices to support neural regeneration, we used porcine derived primary dorsal root ganglion (DRG) sensory neurons which were cultured on non-functionalized and functionalized PPy:Silk scaffolds, and then subjected to pulsed ES at different stimulation potentials ranging from 50-300 mVcm^-1^. The electrically stimulated neurite growth characteristics were evaluated in relation to the scaffolds’ CSC, charge/current injection capacity, and charge-transfer efficiency to identify a potential correlation between them. The effect of enhanced bioactivity and electroconductivity of the scaffolds was also assessed in terms of their capacity to support neural differentiation of primary porcine adipose derived mesenchymal stem cells (pADSCs).

## 2. Materials and methods

### 2.1 Materials

Cocoons of mulberry *Bombyx mori* silk and matured 5^th^ instar larvae of non-mulberry silkworms *Antheraea assama*, were sourced from local sericulture farms, Mangaldai, Guwahati, Assam, India. Pyrrole monomer (Sigma-Aldrich), sodium p-toluenesulfonate (pTS) (Sigma-Aldrich), ferric chloride (FeCl_3_) (SRL), sodium carbonate (Na_2_CO_3_) (Sigma-Aldrich), lithium bromide (LiBr) (Sigma-Aldrich), and sodium dodecyl sulfate (SDS) (Sigma-Aldrich) were used as received.

### 2.2 Regeneration of silk fibroins

Silk fibroin (SF) from *Bombyx mori* (Bm) cocoon was extracted according to the previous protocol.^12, 21^ Briefly, the cocoons were cut and degummed by boiling in 0.02 M Na_2_CO_3_ solution for 15-20 min. The resulting silk fibers were properly washed followed by digestion in 9.3 M LiBr at 60°C for 4 h. The aqueous *Bombyx mori* silk fibroin (BmSF) solution was then subjected to dialysis using 12 kDa cellulose membrane (Sigma-Aldrich) against distilled water for 48 h with regular water changes at every 8 h. The purified silk fibroin solution was then centrifuged at 7500 rpm for 7 min at 4°C to remove any undissolved chunks.

The non-mulberry *Antheraea assama* silk fibroin (AaSF) was isolated from matured 5^th^ instar silkworms following the protocol reported elsewhere.^11^ Briefly, the silkworms were incised, and the posterior silk gland was pinched out. The extracted glands were washed with MilliQ water to remove any traces of sericin. The glands were then then dissolved in 1% (w/v) SDS and dialyzed against ultrapure water at 4°C using 12 kDa dialysis membrane for 4 h. The concentration of both type of the regenerated proteins was determined using gravimetric method and stored in 4°C until further use.

### 2.3 Fabrication of PPy:Silk based aligned scaffold

The degummed Bm fibers were manually arranged in alignment on a microscopic glass slide (25 mm × 75 mm) and firmly secured by attaching adhesive tapes to both longitudinal ends [**Figure 1**]. At least four layers of fibers were employed to establish a consistently aligned platform, achieving a thickness of approximately 1 mm. The aligned fibers were cross-linked by applying a 1% (w/v) silk fibroin solution, followed by an overnight air-drying procedure. It is expected that the reactive amino acid sequences of degummed fibers and the silk fibroin solution will participate in intermolecular interactions through hydrogen bonding or hydrophobic/hydrophilic interactions and will help in binding the aligned fibers together^22^. Afterward, the resultant aligned scaffold was subjected to a 1 h treatment with 70% (v/v) ethanol to trigger β-sheet transition in the silk fibroin coating ensuring its water insolubility and thereby, improves the stability of inter-fiber connections^23^.

Electrically conductive scaffold was prepared by growing polypyrrole nanoparticles (PPy NPs) on the aligned Bm scaffold through *in situ* polymerization of pyrrole using pTS and FeCl_3_ as dopant and oxidant, respectively. Briefly, 14 mM pTS and 38 mM FeCl_3_ were mixed in 30 mL distilled water under stirring for 15 min. The mixture was transferred to a 50 mL falcon tube along with the Bm scaffold, which was kept under shaking condition using a dancing shaker at 4°C for 1 h to allow the dopant and oxidant to be deposited on the fibers. Then, 14 mM pyrrole monomer was added to the falcon tube and again, subjected to continuous shaking for 24 h for polymerization at 4°C. Once polymerization starts, the colour of the mixture turns from yellow to light black. After polymerization, the aligned scaffolds turned into black, resulting in electrically conductive PPy:Silk scaffold. The resultant scaffold was washed thrice with distilled water and air-dried.

The PPy:Silk scaffold was functionalized with 0.5 % (w/v) AaSF solution. The intrinsic RGD tri-peptide present in AaSF is anticipated to enhance the biological functionality of the conductive scaffold. This was accomplished by casting AaSF solution over the PPy:Silk scaffold followed by 2 h incubation at 37°C. Afterwards, the AaSF coating was stabilized by treating the scaffolds with 70% ethanol to induce water insolubility followed. The scaffolds were then washed with distilled water to remove any unabsorbed molecules and air-dried. The PPy:Silk scaffolds before and after functionalization were denoted as PPyBm and AaPPyBm throughout the study. Both types of scaffolds remain stable at room temperature and are stored in airtight packaging for further experimentation.

### 2.4 Physicochemical characterizations

Morphological properties of the fabricated scaffolds were characterized using a field emission scanning electron microscope (FESEM, Carl Zeiss SIGMA, Germany). X-ray diffraction (XRD) studies were performed using an X-ray diffractometer (Rigaku, Model: Micromax-007HF, Japan) in an angular range 10-70° in 2θ, in steps of 0.050°. Chemical composition of the different silk-based scaffolds was investigated with the help of an attenuated total reflectance Fourier transform infrared spectrometer (ATR-FTIR) (PerkinElmer FTIR Spectrum Two, USA) and a micro-Raman spectrometer (LabRAM HR UV-VIS NIR, Horiba). Mechanical properties of the scaffolds were assessed using a Universal Testing Machine (Instron, Model: 5944) equipped with a 100 N load cell at a crosshead rate of 1 mm/min. Steady state current-voltage (I-V) measurement of various mats was tested by a Keithley 2450 source meter using two probe technique at a DC voltage sweep from 0 to ±10 V at room temperature.

### 2.5 Electrochemical studies

Electrochemical performance was investigated using potentiostat/galvanostat (Origalys, OGF01A, France) in a three-electrode cell with the fabricated scaffold (0.5 cm x 0.5 cm) as working, saturated calomel as a reference, and platinum wire as auxiliary electrode. Measurements were conducted in two electrolytes, namely-neurobasal media (NBM) and 1X phosphate buffer saline (PBS). Cyclic voltammetry (CV) analysis was performed at five scan rates of 10, 50, 100, 150, and 200 mVs^-1^ over a potential range of 0.5 to −0.7 V, well within the water electrolysis window. The water window limits are considered between −0.9 to 0.6 V for organic electrodes based on prior studies^6, 7^. The charge storage capacity (CSC) was determined from time integral of current in CV curve.^5, 7^

Electrochemical impedance spectra (EIS) of the various scaffolds was recorded with an AC signal of amplitude10 mV in a frequency range of 0.1 Hz to 100 kHz. The obtained spectra were fitted to suitable equivalent circuit using OrigaMaster 5 software to extract the charge-transfer resistance (R_ct_), solution (electrolytic) resistance (R_s_), and Warburg conductance. Chronoamperometry measurements were carried out with potential steps ranging from 50 to 300 mV and a pulse duration of 1 ms. Chronopotentiometry experiments employed a biphasic oxidation and reduction waveform with equal pulse length and magnitude. These experiments were performed with a pulse duration of 2 ms and applied current steps ranging from 10 to 250 µA.

### 2.6 *In-vitro* degradation study

Biodegradability of the different silk based aligned fibrous scaffolds were assessed in PBS (pH=7.4) with and without protease XIV from Streptomyces griseus (Sigma Aldrich, ≥3.5 U/mg) at 37°C^24^. Protease XIV refers to a non-mammalian enzyme blend employed for in vitro degradation of SF, exhibiting activity specifically in breaking down β-sheet crystalline structures^25^. Hence, it has been documented as the most effective proteolytic enzyme for breaking down SF across a wide range of material formats, such as fibers, films, sponges and hydrogels.

The scaffolds were initially weighed and subsequently, subjected to incubation in PBS with and without 2 U ml^-1^ protease XIV, separately.

At regular 15 day intervals up to 60 days, the scaffolds were rinsed, allowed to air-dry, and then weighed. Throughout the experimental duration, both the PBS and protease solution were replaced every 5 days.

The percentage of remaining mass after incubation was calculated using the following formula:

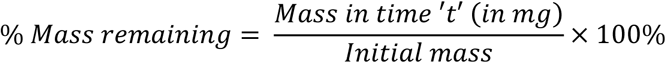

After a 60-day period, the material’s stability was evaluated by examining the altered surface morphology through FESEM and by measuring electrical conductivity via I-V measurements. The films underwent two washes to eliminate salts prior to FESEM analysis. I-V measurements were conducted to ascertain the electrical conductivity under physiological conditions. This study employed three replicates of each sample.

### 2.7 Cell culture

#### 2.7.1 Isolation of dorsal root ganglion (pDRG) sensory neurons

Dorsal root ganglion (pDRG) sensory neurons were also isolated from porcine spinal cord (thoracic and lumbar regions) by adapting previously published protocols with modifications. ^26–29^ Briefly, the spine was rapidly excised post sacrifice (from local abattoir). The spinal cord was carefully detached from the dorsal root and ventral root using standard surgical scissors and fine forceps. Then, the dorsal root ganglion (DRG) was carefully lifted with the help of the roots without touching or damaging the DRG. The extracted DRGs were separated from the roots and cleaned with PBS consisting of 1% penicillin-streptomycin and amphotericin-B (2.5 μg ml^-1^). In this process, 10-16 DRG were obtained at a time. After that, the thin layer of the connective tissue of the epineurium surrounding the ganglion was gently removed. The resultant tissues were subjected to digestion process in 1 % (w/v) collagenase-II in neurobasal media (without any supplement) under gentle agitation for 1 h at 37°C. After digestion, an equal volume of DRG culture media consisting of neurobasal-A medium supplemented with 2% B27, 1% L-glutamine and 1% penicillin-streptomycin and amphotericin-B, was added to neutralize the digestion process. Subsequently, the cell suspension was centrifuged for 5 min at 160 g and the supernatant was removed. The cell pellet was resuspended in complete supplemented neurobasal media and centrifuged again for 5 min at 160 g. The resultant cell pellet was resuspended in 1 mL DRG culture media. Finally, the single cell suspension was plated in Poly-L-lysine coated 60 mm petri-dishes at a density of 6 x10^3^ cells/dish and incubated at 37°C under 5% CO_2_ in air humidified atmosphere. After 1 h, the media was replaced with fresh DRG culture media supplemented with 50 ng/mL NGF. The media was replaced with fresh media to remove non adherent/non-neuronal cells till 3 days from initial plating at an interval of 24 h. After that, one third of the volume of DRG neuronal culture medium freshly supplemented with β-NGF (50 ngml^-1^) is replaced every 3 days. The cells were used for experimentation after two weeks of culture.

#### 2.7.2 Isolation of adipose derived mesenchymal stem cells (pADSCs)

Primary adipose derived mesenchymal stem cells (pADSCs) were isolated from the subcutaneous fat of porcine as reported elsewhere.^30^ In brief, subcutaneous fat from porcine tissue was collected in cold PBS from a local abattoir. The fat tissue was separated from the dermal and muscle portions and subsequently was sliced thinly and rinsed with PBS consisting of 1% penicillin-streptomycin and amphotericin-B (2.5 μg/mL) followed by digestion in 0.1 % (w/v) Type-IA collagenase in incomplete DMEM under gentle agitation for 2 h at 37°C. Equal amount of complete DMEM with 10% (v/v) FBS was added to the digested tissue to neutralize the digestion process. The suspension was subjected to centrifugation twice at 300 g for 15 minutes to separate the stromal vascular portion as pellet. The buoyant adipocytes in the suspension were decanted, and the stromal vascular fraction pellet consisting of the pADSCs was resuspended in DMEM followed by passing the cell suspension through 40 µm cell strainer. The stromal cells were pelleted down by centrifugation and put back in complete low glucose DMEM with 1X antibiotic and amphotericin supplemented with 2 mM L-glutamine and 1 ng/mL basic fibroblast growth factor (bFGF) (Sigma, USA). Lastly, the cells were grown in petri dishes at a density of 5 x10^4^ cells/dish at 37°C under 5% CO_2_. The nonadherent cells in the culture were eliminated by replacing the media after each 24 h till 3 days from initial plating. Passaging was done at 70-90% confluency every 3-4 days. The stromal cells (pADSCs) from passages 3 to 7 were used for experiments.

#### 2.7.3 Neural differentiation of pADSCs

Neural differentiation of ADMSCs was performed according to previous report with slight modification.^35,50,51^ Typically, half of the complete DMEM medium was replaced with initial differentiation media containing DMEM medium supplemented with 10% (v/v) FBS, 1% (v/v) penicillin-streptomycin and 10 ng/mL of bFGF on day 2 and 3. On day 5, the whole media was substituted with complete differentiation medium consisting of 10 ng/mL of each EGF and bFGF along with 1% (v/v) penicillin-streptomycin.

### 2.8 Cell viability assay

The viability of pADSCs and pDRG cells was assessed by measuring their metabolic activity using the alamarBlue™ cell viability reagent (Thermo Fisher) as per the manufacturer’s protocol. Cells were cultured on the different variants of scaffolds (5 mm × 5 mm) in 48 well plates at a concentration of 6×10^3^ cells/well (pADSCs) and 2×10^3^ cells/well (pDRG). After 24, 48, and 72 h, cells were rinsed twice with PBS and then incubated with a 10% v/v alamarBlue reagent for 4 h. The diluted alamarBlue reagents were collected from the culture plate and their fluorescence intensity was recorded at 590 nm. 100% reduced alamarBlue reagent was used as a positive control and the cell culture medium along with scaffold was used as negative control (e.g., Bm+medium, PPyBm+medium and AaPPyBm+medium, are considered as negative controls for their respective scaffold types). Each experiment was conducted in triplicate (n=3).

### 2.9 Live-dead staining

Live-dead staining was carried out to confirm the viability of pADSCs and pDRG cells grown on different scaffolds (Size: 10 mm x 10 mm) using calcein-AM (stains live cells) and ethidium homodimer (EthD-1) (Stains dead cells). Briefly, pADSCs and pDRG cells were cultured on each scaffold, separately, at density of 5 × 10^4^ cells/well and 2×10^3^ cells for 48 h. Then, the cell-seeded scaffolds were washed with 1X PBS twice and incubated at 37°C and 5% CO_2_ with 40 nM calcein-AM and 20 nM EthD-1 for 45 min. The dye solution was removed, and the cell-seeded scaffolds were rinsed again with 1X PBS. Subsequently, the cells were imaged under a fluorescent microscope (EVOS XL digital microscope, Thermo Fisher Scientific, USA) with viable cells appearing green and dead cells appearing red.

### 2.10 Electrical stimulation of pDRG neuronal cells

A bespoke electrical stimulation (ES) set up was made using a 24 well cell culture plate **[Figure S1]**. In brief, the wells were connected with a 0.5 mm platinum (Pt) wire. The wire was placed horizontally on the bottom surfaces of those wells. The different PPy:Silk based conductive scaffolds were then placed over the Pt wire within these wells, guaranteeing direct contact between the scaffolds’ bottom surfaces and the wire. Another Pt wire was inserted vertically into each well, 1 cm apart from the scaffolds/horizontal Pt wire. The vertically positioned Pt wires were partially immersed in the cell culture medium during electrical stimulation of DRG neurons. The set-up was sterilized using 70% ethanol through overnight incubation followed by 1 h UV irradiation inside the biosafety cabinet.

For ES, 2×10^3^ cells were seeded on each of the different PPy:Silk based scaffolds (Size: 10 mm × 10 mm) in the well plate designed for ES. After 24 h of culture, pulsed ES of frequency 50 Hz, pulse width 1 ms with varying amplitude of 50, 100, 200, and 300 mV cm^-1^ was administrated to the cells through different set of scaffolds using an arbitrary function generator (AFG1022, Tektronix) for 2 h each day for 3 consecutive days. Non-stimulated cells grown in the same condition was treated as control. The culture was maintained until Day 7 to study electrically stimulated axonal growth. The experiments were replicated three times, each with a sample size of n=3.

### 2.11 Immunostaining

To affirm the neuronal differentiation of pADSCs, cells were stained against early neuron specific marker-anti-β III tubulin antibody (Abcam, ab18207) and glial cell specific anti-GFAP antibody (Invitrogen, 14-9892-82) as per the manufacturer’s protocol after 14 days of culture on different PPy:Silk based scaffolds. In brief, 5 × 10^4^ cells were seeded on each scaffold (Size: 10 mm × 10 mm) in differentiating medium as described above and incubated for 14 days. For immunostaining, the cell seeded scaffolds were rinsed with 1X PBS thrice and fixed with neutral buffered formalin (NBF). The cells were permeabilized using 0.1% (v/v) Triton X-100 (Sigma Aldrich) in PBS for 15 min. After washing, the cell seeded scaffolds were incubated with 1% bovine serum albumin (BSA) (Sigma Aldrich), 10% goat serum (Sigma Aldrich) and 0.3 M glycine (Sigma-Aldrich) in 0.1% tween PBS (PBST) (Sigma Aldrich) for 1 h. Then, the cell laden scaffolds were treated with 5 μg/mL of primary antibody of interest (anti-β III tubulin or anti-GFAP) in PBST for overnight at 4°C followed by PBS washing and incubation with FITC conjugated secondary antibody Goat Anti-Rabbit (IgG) (Abcam, U.K., 1:2000) for 1 h at room temperature. The cell seeded scaffolds were counterstained with 0.165 μM phalloidin conjugated to rhodamine (Life technologies, USA.) to stain the F-actin and Hoechst-33342 (Sigma Aldrich) for nucleus. The cell laden scaffolds were then visualized under fluorescence microscope (EVOS XL Digital microscope) and representative images are presented. Electrically stimulated and non-stimulated pDRG neurons on different PPy:Silk based scaffolds were immunostained with anti-β III tubulin for axonal growth study using similar method as described above and imaged accordingly.

### 2.12 *In vitro* immunocompatibility assessment

Immunocompatibility of the different silk-based scaffolds was evaluated using murine macrophages RAW 264.7 cells according to the previous protocol^12^. Briefly, 5 × 10^4^ cells were seeded in 24 well plates containing different scaffolds (5 mm x 5 mm) using 500 ng of lipopolysaccharide (LPS) as positive control and non-stimulated tissue culture (NTC) plate as negative control. After 24 h of treatment, conditioned media were collected and tested for nitric oxide (NO) production using a Griess reagent kit, following the manufacturer’s protocol (Invitrogen, USA). The release of interleukin-1β (IL-1β) was monitored using an IL-1β ELISA kit according to the manufacturer’s protocol (Invitrogen, USA). The test was repeated for three times.

### 2.13 Statistical analysis

All tests with a minimum of n = 3 were replicated. One and two-way analysis of variance (ANOVA) analysis (whichever is applicable) were employed to evaluate the statistical significance using GraphPad Prism 10 software. For multiple comparisons, Turkey’s test or Brown-Forsythe test was performed along with ANOVA. Statistical significance was defined at p<0.05. Results are presented as Mean ± Standard deviation (S.D.).

## 3. Results and Discussion

### 3.1 Structural and compositional analysis of biohybrid silk:PPy matrices

Both the structural and compositional characteristics are essential when developing materials for neural applications. The surface roughness can influence cell adhesion and differentiation, whereas the surface topography, such as fiber alignment, can direct the growth of neurites. Likewise, the chemical composition of the surface plays a vital role in determining key factors such as biocompatibility, cytotoxicity, cell adhesion, hydrophilicity, and more. In light of this, before delving deeper into its biological applications, we present an analysis of the structural and compositional features of the developed silk biohybrid, conducted using SEM, XRD, and FTIR techniques. Using our facile fabrication approach, we were able to fabricate aligned Bm silk matrices, which were subsequently functionalised with PPy and AaSF. PPy NPs with avg. dia. of 125 ± 6.67 nm were uniformly deposited through in-situ polymerization technique on the aligned degummed Bm microfibers as can be seen in SEM images [**Figure 2 (a)**]. The results demonstrated that the Bm microfibers (Avg. Dia.= 9.21 ± 1.69 μm) were wrapped with several layers of PPy NPs producing microfibers with an avg. dia. of 12.93 ± 2.53 μm and 13.57 ± 2.22 μm for PPyBm and AaPPyBm, respectively. Bm fibers has reactive functional groups due to the presence of amino acids (such as lysine, tyrosine, serine, glutamate etc.) which help in its chemical modification. Romero et al. systematically showed sulfonic acid modification (also acts precursor for PPy polymerization) of silk promoted pyrrole adsorption on silk substrate resulting in a highly conductive (∼1 S cm^-1^), robust and conjugated PPy:Silk network^14^. A similar prediction was made by Zhao et al., who proposed that hydrophobic interactions between PPy and SF, along with hydrogen bonding and electrostatic interactions, facilitate pyrrole adsorption and subsequent polymerization. Such an interpenetrating strong network of PPy:Silk is resistant to delamination, making it highly significant for use as a hybrid ECB in safe ES protocols. Interestingly, after functionalization with AaSF, the interconnectivity among the PPy NPs became more prominent leading to a smoother surface as compared to PPyBm [**Enlarged view of PS and AaPS in Figure 2 (a)**].

**Figure 2:**
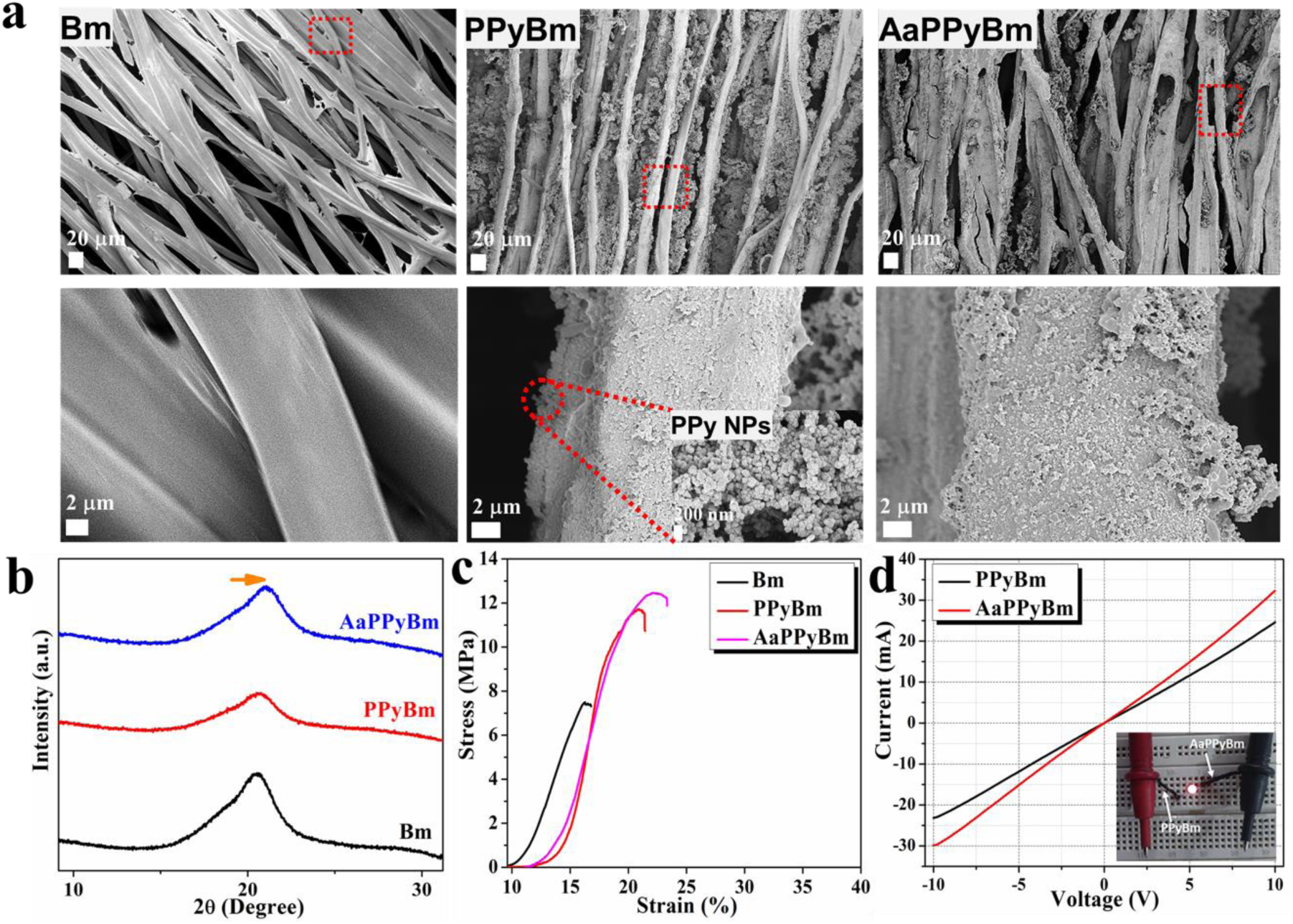
Physico-chemical studies of PPy:Silk based scaffolds. **a.** SEM images of aligned scaffolds of degummed Bm fibers, PPy NPs coated Bm fibers (PPyBm) and Aa silk fibroin (AaSF) functionalized PPyBm (AaPPyBm) as indicated. **b.** XRD patterns; **c.** Representative Stress vs Strain curve and **d.** Room temperature current-voltage (I-V) characteristics of different variants of samples as labeled. Inset of (d) shows illumination of a red LED using PPyBm and AaPPyBm as connector.

XRD analysis revealed a broad but strong diffraction peak at ∼20° present in the diffraction patterns of all scaffolds, corresponding to the β sheet structure both types of silk fibroin [**Figure 1 (b)**]^11^. The results indicated a slight broadening of the diffraction pattern of PPyBm, which is attributed to the amorphous nature of PPy NPs [**Figure S2(A)**]. This is evident from narrowing down the diffraction peak of AaPPyBm with slight shifting towards higher 2θ region, after functionalization with AaSF [**Figure S2(B)**].

FT-IR spectra of the different silk based conductive scaffolds display characteristic fundamental vibrational bands of SF and PPy NPs along with few complex/new peaks as labelled [**Figure 3(a&b)**] and the detailed peak assignment is shown in **Table S2**. FT-IR of aligned Bm fibrous scaffold (treated with 1% regenerated Bm SF followed by 70% ethanol) is mostly dominated by β-sheet structure as evident by appearance of strong peak at 1620 cm^-1^; while a shoulder peak at 1654 cm^-1^, indicates the presence of α-helix/random coil structures^31,32^. In contrast, Aa, (freeze dried from regenerated SF) shows the characteristic amide I peak at 1652 cm^-1^ suggesting its α-helix/random coil conformation. Amide II band in Bm is redshifted (1509 cm^-1^) as compared to Aa (1539 cm^-1^) indicating more β-sheet character (due to increased intermolecular H-boding between different polypeptide chains). The two major fundamental pyrrole ring vibrations appear around 1465 and 1548 cm^-1^ assigned primarily for C=C/C-C stretching mode^33^. The peak at 1548 cm^-1^ has also contributed from N-H bending in secondary amine of PPy^34^. A small peak at 1598 cm^-1^ can be assigned to C=N^+^ vibration of half oxidized pyrrole monomers^34, 35^. The band at 1307 cm^-1^ is attributed to C-N in plane deformation, while the small shoulder near 1359 cm^-1^ can correspond to C-N^+^ stretch with C-C vibration^36^. In addition, several bands appeared with different intensity at 1038, 1088, 1115, and 1161 cm^-1^ are attributed to various C-C-C and C-N-H bending, C-N stretch and N-H deformation as typical aromatic ring vibrations of PPy^36^. The region 1038-1115 cm^-1^ also overlaps with S=O stretching in aryl sulfonate (responsible for compensating the positive charges in PPy) and indicates the doping level in PPy^37, 38^. A small band at 966 cm^-1^ refers to the part of PPy free from the influence of the dopant^39^. The broad signal around 3176 cm^-1^ corresponds to N-H stretching of PPy ring. In PPyBm, amide I band appears with a distinct peak 1622 cm^-1^ (with decreased intensity as Bm fibers are covered by PPy NPs layers of thickness ∼5 μm) with weak signals at 1650, 1682 and 1692 cm^-1^. Past research reported that complex amide I signals of SF in the region at 1640-1650, 1660-1680 and 1670-1695 cm^-1^ are due to deformation/disorderliness, anti-parallel β-sheets and loops of protein’s secondary structure^40^. Hence, the observed signals can be ascribed to probable interaction of PPy with Bm SF, leading to modification in SF polypeptide chains. In the present work, it is to be noted that PPy NPs were grown over the Bm silk fibers during *in-situ* polymerization, to encourage strong coupling between PPy and silk. For that, Bm fibers were first treated with FeCl_3_ and pTSA solution for 10-15 min followed by addition of pyrrole monomer. We assumed that the dopant and oxidant deposition over the fibers would allow polymerization of pyrrole over the silk fibers, creating a PPy layer over those (**Figure 2a**). It is also possible that pTSA can interact with Bm fibers, which provides a strong and stable PPy coating over the fibers^14^. Previously, Xia & Lu showed that cationic radicals of pyrrole monomers during polymerization or fractionally positive PPy chain could be attracted by peptide linkages or carboxylic group of silk, possessing fractional negative charge^41^. These complex events may be responsible for multiple C=O stretching bands suggesting probable covalent/hydrogen bonding/electrostatic interaction between PPy and silk, which is consistent with previous studies^18, 42, 43^. In addition, weak signals around 1690-1720 cm^-1^ might be also an outcome of interaction between terminal carboxylic group of Bm (with secondary amine PPy), resulting in an oxidized PPy^36, 44, 45^. In AaPPyBm, this scenario even becomes more complex with appearance of several bands, particularly at 1617, 1663, and 1698 cm^-1^ [**Figure 3 (b)**]. It is evident that relatively weak band at 1617 cm^-1^ (red shifted) and the strong band at 1663 cm^-1^ (blue shifted) have contributions from amide I in PPyBm and Aa. Interestingly, the small but distinct band at 1698 cm-1 clearly suggests the C=O stretching due to-COOH group (of Aa) conjugated with secondary amine of PPy^46^. Previous studies showed that deprotonation of PPy, when treated with carbonyl group containing element and deprotonated/oxidized PPy shows a signature peak in the region 1690-1730 cm^-1^ ^44, 45^. Hence, the small bands in the region 1690-1737 cm^-1^ indicate a highly oxidized form of PPy due to covalent/hydrogen bonding/electrostatic interaction between PPy and Aa. One of the interesting observations to support this claim is the disappearance/blue shifting of N-H stretching vibration band (which appeared with a small signal in PPyBm at 3278 cm^-1^), towards ∼3400 cm^-1^, overlapped with O-H stretching vibration of silk fibroin (seen in case of both types of silk). In fact, the strong bipolaronic band of pure PPy at 1161 cm^-1^, shifted to 1154 and 1160 cm^-1^ in PPyBm and AaPPyBm, which also indicates possible π-π interaction between PPy and silk fibroin^47^.

**Figure 3:**
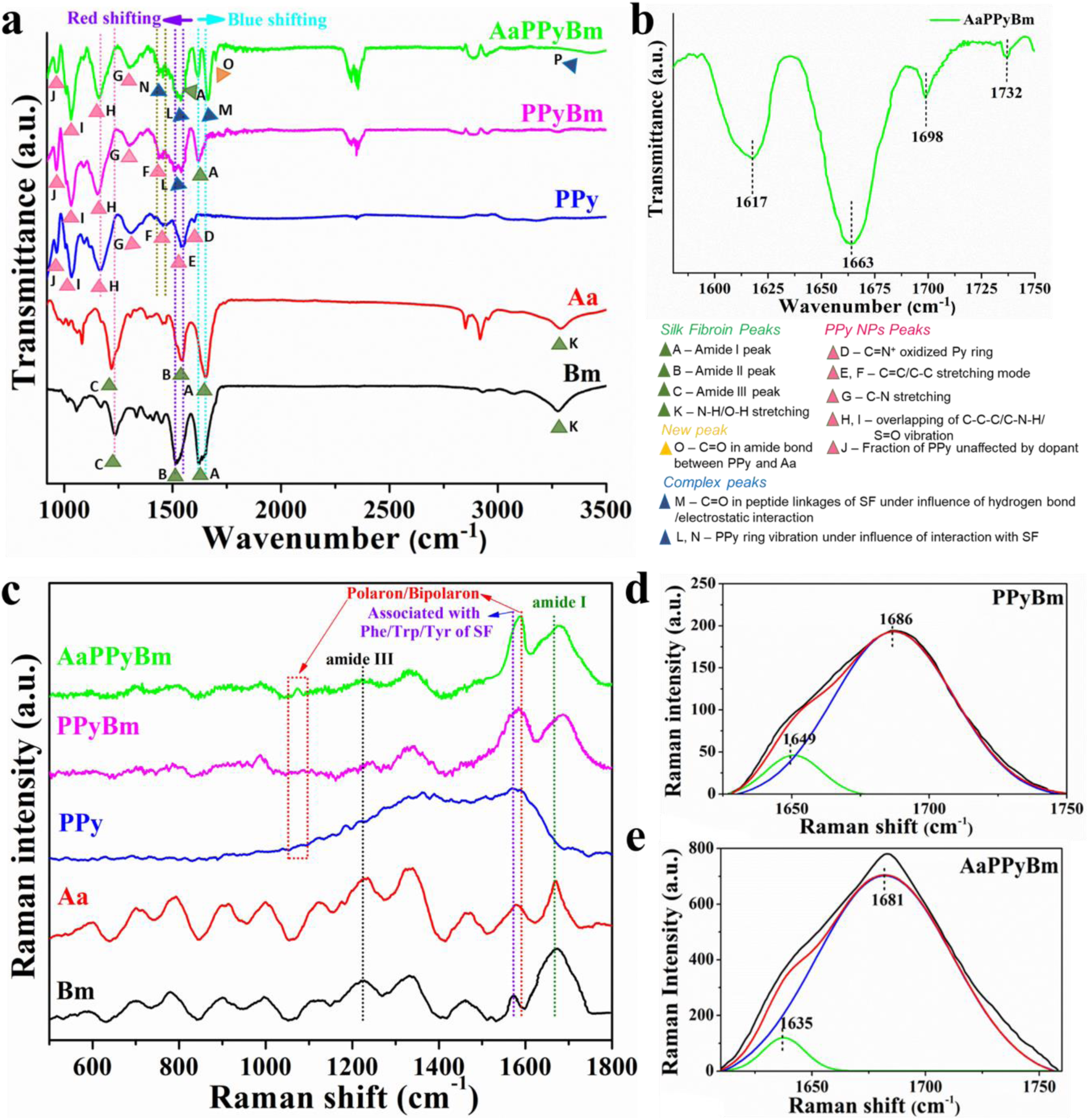
Chemical compositional analysis results showing FT-IR patterns (a & b) and Raman spectra (c-e). FT-IR spectra of (a) various silk-based scaffolds including pristine PPy NPs as labeled and (b) AaPPyBm highlighting the major peaks. Raman spectra (c) various PPy:Silk based scaffolds along with pristine Aa, Bm and PPy NPs. Lorentzian deconvolution and curve-fitting of amide I peak of (d) PPyBm and (e) AaPPyBm.

Raman spectroscopic analysis recorded with 532 nm excitation wavelength, also reveals fundamental vibrational bands of both SF and PPy NPs along with few complex signatures, indicating probable interaction among the different constituents [**Figure 3(c-e)**]. Consistent with the FT-IR results, Raman analysis also suggests the dominance of β-sheet structure in pure Bm scaffold with strong amide I and amide III signals at 1675 and 1227 cm^-1^ ^48^. In Aa, the amide I peak is redshifted to 1665 cm^-1^, which is comparatively narrower than that of Bm, indicating the presence of α-helix/random coil conformation, while amide III peak appears at 1237 cm^-1^. The extended alanine sequence and polyalanine, which impart enhanced mechanical resilience to Aa as compared to Bm, gives rise to stronger signature peaks at 901-916 and 1337 cm^-1^, respectively^48, 49^, as compared to those appeared at 899 and 1334 cm^-1^ in Bm. Pure PPy NPs exhibit mainly two strong broad bands with multiplets in the region 1327-1390 and 1570-1592 cm^-1^. The strong broad peak in the region1570-1592 cm^-1^ corresponds to the backbone stretching of C=C mode and inter-ring C-C mode in the backbone of the polaron structure^50–52^. In fact, this region is useful to identify the neutral, oxidized (polaronic), and fully oxidized (bipolaronic) state of PPy, corresponding to the bands at 1560, 1580 and 1610 cm^-1^, respectively. The weak bands in the region 1458-1502 cm^-1^ relates to the skeletal, or C-C and C-N, stretching vibrations^53^. The peak at 1327 cm^-1^ is attributed to the neutral state of PPy, while the two distinct multiplets at 1366-1390 cm^-1^ are assigned to the antisymmetrical inter-ring stretching C-N vibration of oxidized (doped) PPy^54, 55^. The weak signals around 1186-1270 cm^-1^ are assigned to the different C-H in plane bending vibrations in pyrrole ring^53, 55^. Another important signature of conductive PPy is N-H in plane vibration due to polaronic/bipolaronic state^51, 52^, which feebly appears at 1075 cm^-1^. This particular vibration is highly sensitive to the presence of an anion in the protonating acid^56^.

Similar to FT-IR analysis, the Raman active bands due to SF are diminished in PPyBm due to growth of PPy NPs layer over the Bm fibers, with shifting of several bands. Most importantly, in AaPPyBm, the strong and sharp band at 1595 cm^-1^ corresponds to the fully oxidized bipolaronic state of PPy, which is also supported by the distinct peak at 1074 cm^-1^ due to N-H in plane signature peak for deprotonated PPy^52, 56^. It indicates that anionic AaSF can act as a dopant by inducing deprotonation at secondary amine site of the PPy ring, which causes shifting of the bipolaronic band towards high frequency region. This is also consistent with prior studies^50, 51, 56^. The possible interaction of Aa with PPy is also indicated by shifting of amide I peak from 1686 cm^-1^ (in PPyBm) to 1681 cm^-1^ in AaPPyBm with a clear shoulder peak 1630-1650 cm^-1^. To further elucidate the probable interaction mechanism, amide I peak of PPyBm and AaPPyBm was subjected to Lorentzian deconvolution and curve-fitting [**Figure 3(d&e)**]^57–59^. Curve fitting resulted in two bands in PPyBm (1649 and 1686 cm^-1^) and AaPPyBm (1635 and 1681 cm^-1^). C=O stretching is mainly responsible for amide I band of SF and its stretching vibration is influenced when there is an interaction, like reactive secondary amines of PPy. Both PPyBm and AaPPyBm were subjected to ethanol treatment for β sheet induction, which is supported by the deconvoluted amide I band at 1686 and 1681 cm^-1^, respectively and in agreement with the FT-IR results. The band observed at 1649 and 1635 cm^-1^ in PPyBm and AaPPyBm, respectively, can be attributed due to the C=O stretching in aromatic side chains^60^. Previous study described such amide I specific Raman vibrational bands at low wavenumber region owing to the conjugated C=O of tertiary amide^61^. It suggests probable interaction of amino group of PPy and terminal carboxylic group of Bm or Aa. The intensity of the peak corresponding to this interaction appears to be stronger in AaPPyBm than that of PPyBm as revealed by the peak deconvolution and curve fitting [**Figure 3(d & e)**]. This also supports highly oxidized form of PPy in AaPPyBm, leading to its enhanced electrical conductivity, which also aligns with the observation revealed by FT-IR analysis.

### 3.2 Mechanical behaviour of biohybrid silk:PPy matrices

Owing to the unique mechanical properties of nerve tissue in terms of elasticity and flexibility, it is essential to design neural scaffolds with similar mechanical characteristics to provide an appropriate microenvironment for nerve regeneration^62^. A highly rigid or non-elastic biomaterial scaffold can induce stress concentration, inflammation, and damage to the surrounding tissue, impeding the regenerative process. Elasticity of biomaterial scaffolds were shown to effect morphology and functions of nerve cells^63^ as well as neural stem cells^64^. In addition, Rosso *et al.* highlighted the mechanoresponsive behaviour of Schwann cells and dorsal root ganglia in relation to the substrate stiffness^65^. Hence, the mechanical strength test of the fabricated aligned scaffolds was conducted to confirm their potential implication as nerve guidance channel (NGC).

Stress *vs* Strain behavior of different aligned scaffolds and important parameters derived from it are shown in **Figure 1 (c)** and **Table 1**, respectively. The results show that the incorporation of PPy NPs onto the Bm microfibers enhanced the mechanical properties of the scaffold significantly which was further enhanced by AaSF functionalization. AaPPyBm exhibited highest stiffness constant followed PPyBm and Bm. Further, the ultimate tensile strength (UTS) and elongation at break (%) were also enhanced in the order of AaPPyBm>PPyBm>Bm [**Table 1**]. The enhanced mechanical properties of AaPPyBm may be due to the induced interconnectivity/cross-linking among the PPy NPs after AaSF functionalization, as suggested by SEM analysis. Studies showed that human tibial nerve has elastic moduli in the range of 8-16 MPa, and UTS values in the range of 6.5-8.5 MPa^66^. This suggests that the fabricated aligned scaffolds are mechanically stronger than the native human nerve, which is essential for their potential implications as NGC. It is because the implantable NGCs should be capable of withstanding physiological load as well as should be stiff enough to support suturing and electrodes for external electrical stimulation^67^.

**Table 1:** Mechanical properties of Bm, PPyBm and AaPPyBm.

| Sample Name | Elastic modulus<br>(MPa) | Ultimate<br>tensile<br>strength<br>(MPa) | Elongation at<br>break<br>(%) |
| --- | --- | --- | --- |
| Bm | 162.39±11.30 | 7.49±1.12 | 16.27±3.07 |
| PPyBm | 289.76±4.81 | 11.71±1.47 | 20.90±2.82 |
| AaPPyBm | 293.45±7.14 | 12.47±2.23 | 21.67±4.30 |

### 3.3 In-vitro biodegradation and stability study

Matrix remodeling is essential to facilitate neurite extension as well as to support neural progenitor stemness and differentiation.^68, 69^ We simulated the proteolytic degradation by incubating the scaffolds in proteases to assess the biodegradability of the developed matrices. When silk-based biomaterials undergo proteolytic degradation, it results in the fragmentation of silk fibroin (SF) into smaller polypeptides, eventually breaking down into amino acids that can be readily absorbed or metabolized within the body^25^ serving as a bioresorbable scaffolding matrix. It is evident from the mass degradation profile of Bm scaffold, which showed ∼80% mass loss in proteolytic condition after 60 days as compared to ∼35% mass loss in only PBS [**Figure 6(A&B)**]. However, the degradation behavior in both enzymatic condition and PBS were slowed down, when the Bm fibers are coated with PPy NPs. Around 40-50% mass loss was observed for both PPyBm and AaPPyBm under proteolytic action against 20-25% mass loss in only PBS [**Figure 6(A&B)**]. This observation is substantiated by the FESEM images of the degraded scaffolds, which clearly reveal fiber fragmentation due to proteolytic activity and the partial removal of the PPy coating in both PPyBm and AaPPyBm, in contrast to the relatively stable and intact surface morphology observed in PBS control (without protease) [**Figure 6(C&D)**]. The representative photographs of different degraded scaffolds are shown in **Figure S9**. It’s noteworthy that PPyBm and AaPPyBm maintained their conductive properties even after being incubated in both the PBS control and protease solution for a duration of 60 days, falling within the range characteristic of ideal semiconductors [**Table 2 & Figure S10**]. The findings indicate that PPy NPs play a crucial role in preserving the electrical and structural stability of the scaffolds in the presence of proteolytic activity. This underscores the potential for adjusting the biodegradability of these conductive scaffolds. Such a property holds relevance in the context of nerve regeneration, which typically demands a long-term stability of biomaterials owing to their slower growth rate as compared to other tissues.

**Table 2:** Electrical conductivity of PPyBm and AaPPyBm.

| Sample Name | Conductivity (S cm<sup>-1</sup>) |  |  |
| --- | --- | --- | --- |
|  | (as fabricated) | After 45/60 days of incubation |  |
|  |  | PBS (under hydrolytic conditions) | Under proteolytic conditions |
| PPyBm | $6.23 \times 10^{-2} \pm 0.98$ | $1.49 \times 10^{-3} \pm 0.98$ | $5.53 \times 10^{-4} \pm 1.03$ |
| AaPPyBm | $9.18 \times 10^{-2} \pm 1.19$ | $1.70 \times 10^{-3} \pm 0.80$ | $3.56 \times 10^{-4} \pm 0.56$ |

### 3.4 Electrical conductivity studies

Electrical characterization by current-voltage characteristics measurement of the PPy:Silk based scaffolds with and without AaSF functionalization demonstrated their highly conductive nature with conductivity of 6.23 and 9.18 mS cm^-1^ for PPyBm and AaPPyBm, respectively [**Figure 1(d) & Table 2**]. Interestingly, AaSF functionalization was found to contribute towards increased conductivity of AaPPyBm. Previous studies used a plethora of negatively charged bioactive molecules such as hyaluronic acid (HA), chondroitin sulphate (CS)^70^, lysine^71^, glutamate^72^, RGD peptide^73^, laminin and fibronectin like fragments^74^, etc. as anionic dopants in the synthesis of biologically functionalized PPy, which also controls its physical and chemical properties. Similar to these bioactive molecules, AaSF also possesses negative Zeta potential at pH=6.9 [**Figure S3(A)**], which is due to the higher oxygen containing functionalities (e.g., carboxylic groups) than the amino groups. PPy NPs synthesized with FeCl_3_ as oxidant (without using any dopant) demonstrates positive surface potential (after purification) [**Figure S3(B)**]. This is due to the positively charged nitrogen atoms as shown by Zhang *et. al* ^75^. However, purified PPy NPs synthesized with FeCl_3_ as oxidant and pTS as dopant again exhibited negative Zeta potential at the same pH, owing to acidic functional groups such as SO_3_^-^ and Cl^-^ [**Figure S3(C)**]. Interestingly, when pTS was replaced with AaSF, the resultant PPy also demonstrated slightly negative zeta potential, indicating the anionic dopant like role of AaSF [**Figure S3(D)**]. This is further supported highest negative Zeta potential of AaSF modified PPy NPs synthesized using FeCl_3_ and pTS [**Figure S3(E)**]. Consequently, AaPPyBm showed enhanced conductivity than that of PPyBm. To further support this claim, two different variants of PPyBm were synthesized and their conductivity was measured [**Figure S4**]. First, PPyBm fabricated with FeCl_3_ as oxidant and without using any dopant. The other variant was prepared by replacing pTS dopant with AaSF. Interestingly, the latter showed greater conductivity than the former. This suggests that AaSF functionalization not only provides the highly interconnected network PPy NPs leading to increased mechanical behaviour of PPy:Silk scaffolds, but also contributed towards its enhanced electrical conductive property.

### 3.5 Electrochemical studies

Characterization of the electrochemical characteristics of scaffolds under biologically relevant conditions has been shown to evaluate properties such as charge storage, charge/current injection capacity, and charge-transfer kinetics between the electrode (scaffold)-tissue/cell interface^6, 76–78^. Hence, electrochemical behavior was recorded in three electrode configurations with various silk based electroconductive scaffolds as working electrodes in two biologically relevant electrolytes, namely, 1X PBS (pH=7.4) and neurobasal media (NBM). Cyclic voltammetry (CV) profiles of all the scaffolds were recorded in both the electrolytes at scan rates of 10, 50, 100, 150 and 200 mVs^-1^ within a potential range of −700 mV to +500 mV [**Figure 4(A, B, D & E)**]. The obtained voltammograms of PPyBm and AaPPyBm exhibit variations in their patterns in both electrolytes, highlighting distinctions in their electroactivity derived from differences in redox states or charge states. The electrochemical behaviour of the scaffolds depicting their redox activity both in NBM and PBS are summarized with their redox potentials and peak currents (anodic & cathodic) in **Table S3 & S4**. The data depicts an increase in the redox peak potentials for each scaffold in both the electrolytes with increasing scan rates (u), while the redox peak currents vary linearly with u^1/2^ [**Figure S5 & S6**]. Such characteristics indicate the quasi-reversible behavior of PPyBm and AaPPyBm, where peak current and scan rate (u) are dependent on electron transfer rate constant^79–81^.

**Figure 4:**
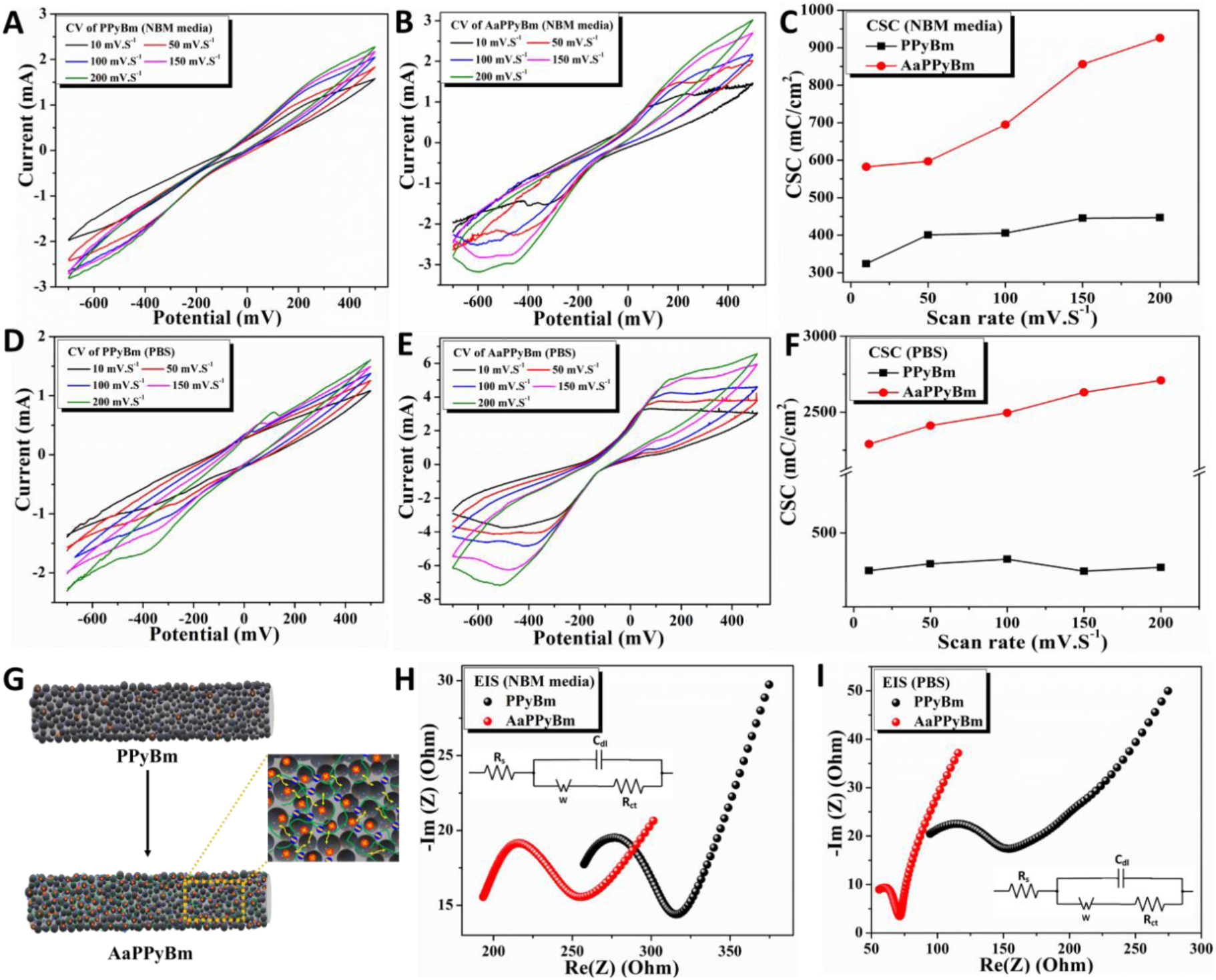
Electrochemical behavior of various electrodes in a three-electrode configuration assessed in neurobasal media (NBM) and 1X phosphate buffer saline. Cyclic voltammetry of PPyBm and AaPPyBm in NBM **(A&B)** and PBS **(D&E)** at scan rate of 10, 50, 100, 150 and 200 mVs^-1^. Charge storage capacity (CSC) of PPyBm and AaPPyBm in **(C)** NBM and **(F)** PBS at different scan rates. **(G)** Schematic illustration showing enhanced ion diffusion/movement of charge carriers (indicated in yellow arrows) in AaPPyBm due to anionic AaSF (indicated in green) connecting PPy NPs (indicated in black). Nyquist plots showing electrochemical impedance spectra (EIS) of PPyBm and AaPPyBm in **(H)** NBM and **(I)** PBS in frequency range from 0.1 Hz to 100 kHz. Insets of **(H)** and **(I)** display equivalent circuit used for fitting to determine the parameters shown in **Table 3**.

**Table 3:**
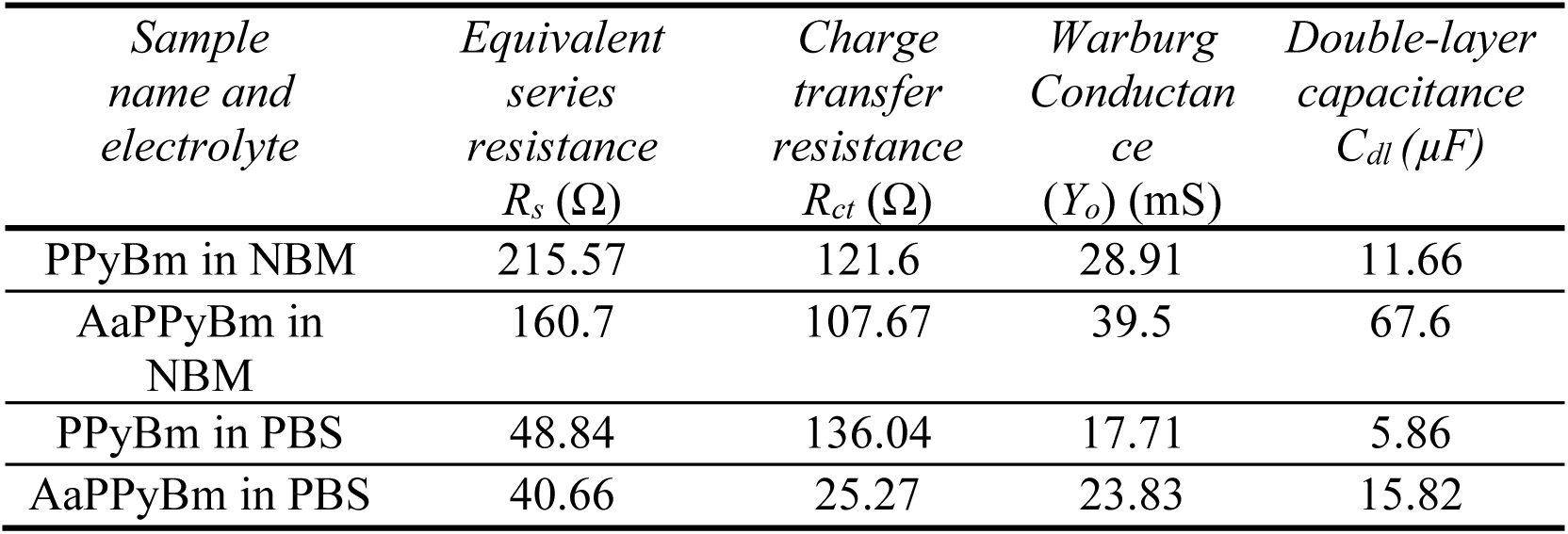
Equivalent circuit fitting parameters for different scaffold materials in NBM and PBS, derived from the corresponding Nyquist plots shown in. **Figure 4(H&I).**

In NBM, PPyBm displays relatively a weaker anodic and cathodic peak, whereas in PBS, these peaks are more distinct [**Figure 4(A&D)**]. In both NBM and PBS, the anodic and cathodic peaks shift towards more anodic and cathodic direction, respectively, with increasing scan rate (u). An anodic peak represents anion insertion into PPy, while a cathodic peak is associated with a cation insertion^79, 82, 83^. In PBS, both of these peaks are more pronounced indicating better redox activity of PPyBm compared to that in NBM. This is further supported by better linearity (Adj. R-Square ∼ 1) of the correlation between peak current(s) and u^1/2^ in PBS, indicating the improved redox reversibility and faster charge transfer, when compared to that in NBM [**Figure S5**].

On the other hand, the CV profiles of AaPPyBm in both the electrolytes display a distinct broad anodic peak and two cathodic peaks with greater area under the voltammograms [**Figure 4(B&E)**]. The two clear cathodic peaks are indicative of higher reduction tendency (cation insertion) the scaffold. This can be interlinked with the fully oxidized (bipolaronic) form of PPy induced after functionalization with Aa, as suggested by FT-IR and Raman analysis [**Figure 3**]. In fact, the reduction of AaPPyBm becomes more pronounced with an increased area as the scan rate (u), suggesting the ability of the scaffold to accommodate more cation from the electrolyte with higher charge transfer efficiency. This is consistent with the enhanced linearity (Adj. R-Square ∼ 1) of the correlation between peak current(s) and u^1/2^ in both NBM and PBS for AaPPyBm than that of PPyBm [**Figure S6**]. The observation suggests rapid charge transfer between AaPPyBm and the electrolyte(s) as well as the reversibility of the redox processes. The cyclic voltammograms were further utilized to determine the charge storage capacity (CSC) of the scaffolds [**Figure 4(C&F)**]. AaPPyBm exhibits significantly higher CSC values in both the electrolytes as compared to PPyBm. Particularly, AaPPyBm showed a maximum CSC of 925 mCcm^-2^ at 200 mVs^-1^ in NBM and 2708 mCcm^-2^ at 200 mVs^-1^ in PBS. Both PPyBm and AaPPyBm possess very high CSC values, when compared to bionic devices or conventional electrodes such as iridium oxide (∼4-127 mCcm^-2^)^84, 85^, titanium nitride (∼22-253 mCcm^-2^)^86, 87^, platinum or platinum iridium alloys (∼16-22 mCcm^-2^)^88^.

Electrochemical impedance spectroscopy (EIS) measurement of the conductive scaffolds was performed in NBM and PBS to gain a deeper insight about the electrochemical charge-transfer kinetics in terms of resistance and diffusion characteristics over a frequency range of 0.1 Hz to 100 kHz. The data were plotted with the imaginary impedance (Z ̋) versus the real impedance (Z’) at each sampled frequency giving the Nyquist plot [**Figure 4 (H&I)]**. In ideal condition, a perfect capacitor displays a vertical line with only the imaginary impedance (Z ̋) component. The Nyquist plots of PPyBm and AaPPyBm in both the electrolytes depict a semi-circle in the high-frequency region, followed by a straight line in the low-frequency range, indicating the deviation from the ideal capacitive characteristics. To further understand the reaction kinetics in the overall electrochemical activity of the silk based electroactive scaffolds, these data were fitted to an equivalent circuit model using OrigaMaster 5 software [**Inset(s) of Figure 4(H&I)**], where each circuit element is selected to correspond to a real physical component in the electrochemical system^89^ and the fitting parameters are shown in **Table 3**. The X-axis intercept of the Nyquist plot in the high-frequency range represents the electrolyte resistance and termed as series or solution resistance (R_s_) in the equivalent circuit. The solution resistance (R_s_) is primarily influenced by the physical characteristics of cations/anions of the electrolyte and the electronic state of the electrode (i.e., scaffold). Studies reveal that cations/anions with smaller ionic/hydrated sphere radius promote ion adsorption at electrode surface, while the greater ionic mobility and conductivity boost charge transfer process.^90, 91^ In the present study, AaPPyBm exhibits different R_s_ values in both the electrolytes, which are lower than PPyBm [**Table 3**]. As discussed above, AaPPyBm has more electrically active PPy due to generation of bipolarons after anionic Aa functionalization and is more susceptible for interactions with the cations of electrolytes, as compared to PPyBm [**Figure 4 (G)**]. NBM has several salts that can produce cations such as Na^+^, K^+^, Ca^2+^, Mg^2+^ and Fe^3+^ during electrochemical reaction [**Table S4 and S6**], and the electronic nature of AaPPyBm is expected to promote cation adsorption with more ease, leading to a significantly lower R_s_ in NBM^92, 93^. This is consistent with the lower Rs value in 1X PBS as revealed by AaPPyBm. However, the difference is less as PBS has relatively less concentration of cations as compared to NBM [**Table S5 and S6**]. As stated, the semi-circle in the high to mid frequency range suggests a non-ideal capacitive system with leakage channels and the electrochemical interactions in this frequency range can be used to get valuable information about charge-transfer processes. Hence, this is represented in terms of a double-layer capacitance (C_dl_) at the electrode surface due to the electrolyte ions, in parallel connection with a resistor in the equivalent circuit. The resistor is presented as charge-transfer resistance (R_ct_) that relates to the kinetics of the heterogeneous electrochemical processes, implying the resistance encountered by the ions/charges at the electrode (i.e., scaffold)-electrolyte interface during redox reactions. The diameter of the semi-circle represents R_ct_ values. The results show significantly lower R_ct_ values for AaPPyBm both in NBM (107.67 Ω) and PBS (25.27 Ω) than that of PPyBm [**Table 3**]. This is also evidenced by the higher current values observed in the CV profiles and increased conductivity (9.18 mS cm^-1^) for AaPPyBm. This is again related to the possible role of AaSF as an anionic dopant, which converts partially oxidized PPy to fully oxidized (bipolaronic) PPy. Thus, the resultant high electroactivity of AaPPyBm eases the movements of charge carriers across the electrode-electrolyte interface. The impedance response profile, in the low-frequency range, also includes another important parameter due to the resistance encountered by the electrolyte ions during diffusion within the electrode (i.e., scaffold), termed as Warburg resistance (W) in the equivalent circuit and listed in the form of Warburg conductance (Y_o_) [**Table 3]**. Fitting parameters reveal that enhanced Warburg conductance (Y_o_) for AaPPyBm in both the electrolytes, indicating the lower ion diffusion resistance as compared to PPyBm. The findings indicate that the incorporation of Aa silk fibroin into PPyBm serves to expedite charge transfer at the electrode-electrolyte interface [**Figure 4(G)**]. Additionally, it facilitates ion diffusion within the scaffold with ease, leading to improved electrochemical kinetics of the electrode.

Electroactive scaffolds intended for use in electrically stimulated tissue repair, should be also electrochemically stable. The charge-transfer reactions occurring at the electrode ( scaffold)-electrolyte interface, whether they are capacitive or faradaic, should not have detrimental effect on the scaffolds through their degradation (under constant electrical pulsing) or on the surrounding tissues^6, 94^. Both PPyBm and AaPPyBm demonstrate good electrochemical stability, when they were tested for cyclic stability using CV measurements upto 200 cycles at 200 mVs^-1^ in NBM [**Figure S7**]. Voltammograms of both the scaffolds display an increase in the enclosed area under the curves with increasing cycle numbers. During the repeated cycling, the electrolyte ions penetrate more into the interior of the scaffolds, and thereby activate the additional active sites during the process of cycling. Consequently, it can increase the overall capacitance of the scaffolds, which is manifested from the larger CV area of the voltammograms with increasing cycle number.

Neuroprosthetic or neuromodulation devices employ either constant voltage or constant current pulses to deliver therapeutic electrical stimulation. This approach is utilized in treating deafness through cochlear stimulation as well as addressing neurological diseases and disorders through deep brain stimulation and spinal cord stimulation^95–98^. It is essential to carefully select the stimulation parameters well within the threshold for electrode polarization and tissue damage, which are traditionally described by the Shannon equation^99^. It can be estimated from the charge and current injection capacity of an electrode under a bias voltage or potential range, which is within the water electrolysis widow limit (reduction and oxidation of water). Clinical applications involving therapeutic electrical stimulation utilize a biphasic charge-balanced waveform for charge injection^100, 101^. To determine the charge and current injection capacity of the various silk based electroactive scaffolds, chronoamperometric technique was employed to record the current responses in NBM under a constant charge balanced voltage waveform with a bias voltage ranging from 50 to 300 mV and a pulse width 1 ms [**Figure 5 (A&B)**]. The bias voltage range is well within the water electrolysis potential window, which was previously shown to be typically from −0.6 V to 0.8 V (−0.9V to 0.9V) versus Ag/AgCl^7^. 1 ms pulse width was chosen as this was the shortest pulse achievable with the potentiostat/galvanostat (Origalys, OGF01A, France) and similar pulse width was also utilized for stimulation of DRGs using a conducting polymer platform^102^. Both oxidative and reductive current values increase with rising bias voltage for both PPyBm and AaPPyBm. However, at a given bias voltage (except at 50 mV), AaPPyBm exhibits higher oxidative and reductive current values compared to PPyBm. It suggests improved charge injection processes in AaPPyBm, which is also in agreement with increased redox activities as observed in CV analysis [**Figure 4 (B&E)**]. The time integral of the biphasic (cathodal and anodal) current pulse at a bias voltage gives the charge injection capacity of the electrode and the results are shown in **Figure 5(E)**. AaPPyBm demonstrates significantly higher charge injection capacity at all experimental bias voltages, when compared to that of PPyBm. Interestingly, AaPPyBm has comparable or higher charge injection capacity even at low potential or at high potential (e.g., 0.46 mC cm^-2^ at 50 mV and 3.52 mC cm^-2^ at 300 mV), when compared to that of conventional/traditional bioelectrodes such as Titanium nitride (TiN) (0.9 mC cm^-2^)^85^, platinum/platinum-iridium alloy (100-350 μC cm^-2^)^103^, sputtered iridium oxide films (SIROFs) (750 μC cm^-2^)^104^, PPy/Pt (1.3 mC cm^-2^)^7^, PEDOT/ITO (3.6 mC cm^-2^)^105^, and activated iridium oxide flms (AIROF) (5 mC cm^-2^)^6^. Accordingly, AaPPyBm also shows significantly higher current injection capacity than PPyBm [**Figure 5(F)**]. Studies reported the charge-density threshold for various neural electrodes are in the range from 5-4000 μC.cm^-2^ ^106, 107^. Further, chronopotentiometry technique was also utilized to record the voltage response of PPyBm and AaPPyBm in NBM at current density from 10-250 μA cm^-2^ at pulse width of 2 ms [**Figure 5(C&D)**]. The results show that both the electrodes depict symmetric and stable potential profiles within the applied current density range affirming their sustainability and stability. The voltage derivative plots and their reciprocal also confirm redox stability of the electrodes [**Figure S8(A&B)**], which aid in understanding the electrochemical reaction mechanisms at the electrode interface^108^. Ideally, any capacitance current would appear as constant and any faradaic current as sharp dip. Also, the reciprocal of the derivative would transform any faradaic current into sharp peak, which can be seen both in case of PPyBm and AaPPyBm [**Figure S8(A&B)**]. The results show both capacitive and faradaic reactions are responsible for the electrochemical behaviour of the scaffolds. Notably, capacitive currents dominate at low current density, while it can be visualized the ratio of faradaic to capacitive current increases at higher current density. The findings reveal that the current silk based electroactive electrode, particularly AaPPyBm has the potential to deliver therapeutic electrical stimulation at a low electrical potential without causing any electrode (i.e., scaffold) degradation and tissue damage.

**Figure 5:**
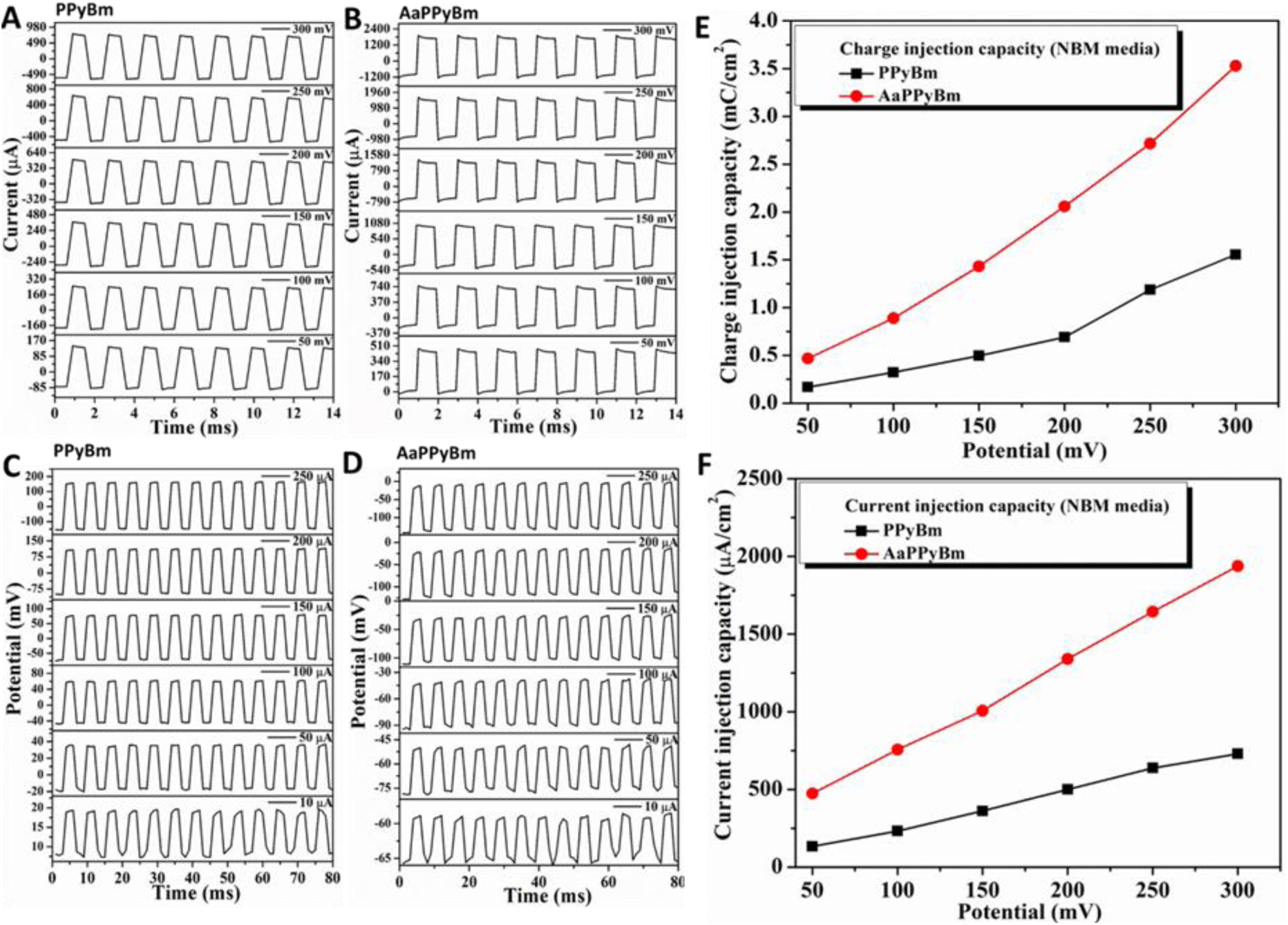
Chronoamperometry patterns of **(A)** PPyBm and **(B)** AaPPyBm showing current response at different stimulation potentials in NBM. Multiple pulse chronopotentiometry of **(C)** PPyBm and **(D)** AaPPyBm depicting voltage transients in response to injected current limits as indicated. **(E)** Charge injection capacity and **(F)** current injection capacity of the conductive scaffolds derived from their chronoamperometric responses at various potentials as indicated.

### 3.6 *In-vitro* biocompatibility study

Cytocompatibility of the various silk-based scaffolds was tested in terms of %reduced Alamar Blue reagent by the pADSCs and pDRGs seeded on those scaffolds after 24, 48 and 72 h [**Figure 7(A&B)**] and normalized to tissue culture plate as controls. Both cell types demonstrated increased % Alamar Blue dye reduction with increasing duration of culture time on all the scaffolds, suggesting increased cellular metabolic activity. The results indicate that all the silk-based scaffolds are cytocompatible and support cellular growth. This is consistent with the well-explored biocompatibility of both silk fibroin and PPy.^3, 8, 10, 20, 109^ The results further show that pADSCs cultured on AaPPyBm demonstrated significantly higher % reduction, when compared to that on pure Bm scaffolds at culture time of 48 (p≤0.01) and 72 h (p≤0.001) [Figure..A]. pDRGs seeded on the AaPPyBm also showed relatively higher reduction of alamarBlue, however, they are not statistically different [**Figure 7(B)**]. pDRG consists of heterogenous populations of sensory neurons and mostly acquire a non-proliferative state unless stimulated. The seeded pDRG cells remained viable but did not exhibit proliferative potential similar to the pADSCs. Moreover, the better cell viability on AaPPyBm scaffolds can be attributed to the presence of cell affinitive RGD (arginine-glycine-aspartate) tripeptide in AaSF leading to the improved cell-biomaterial interactions. The confirmation of cell viability on the various silk-based scaffolds, as previously discussed through the reduction of alamar blue, was further validated by conducting live-dead staining of pADSCs and pDRGs after a 2-day culture period [**Figure 7(C&D)**].

**Figure 6:**
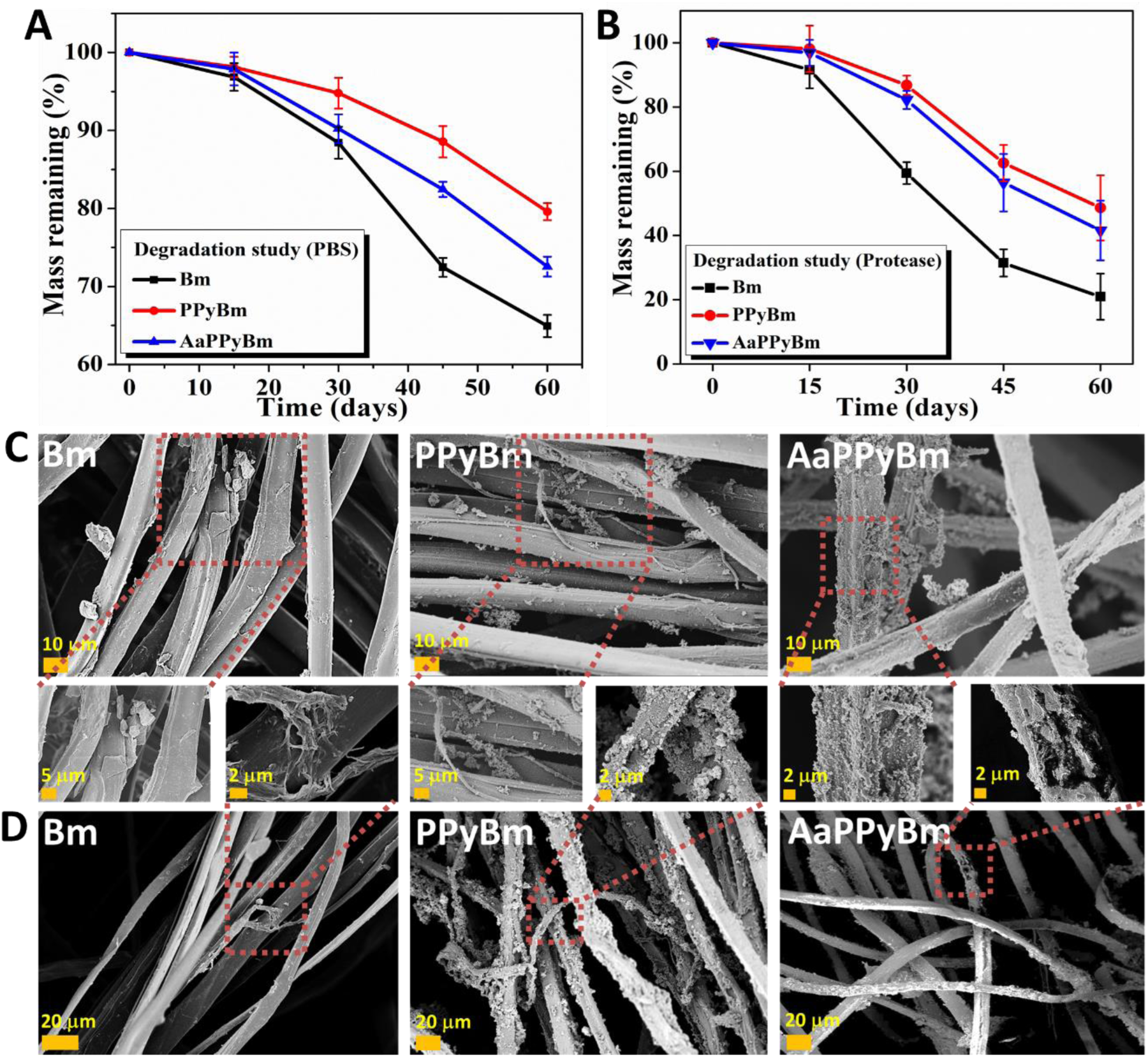
*In-vitro* biodegradation results. Percentage of remaining mass profiles in **(A)** PBS (pH=7.4) and **(B)** protease XIV solution of Bm, PPyBm and AaPPyBm after 60 days. Representative FESEM images of various degraded scaffolds incubated in **(C)** PBS and **(D)** protease solution as indicated. Magnified images are highlighted in red dotted squares of different scaffolds to show the impact of degradation in PBS and protease. Scale bar = 10 μm (**C**), 20 μm and 5 μm (Magnified images); Data in (**A) & (B)** were expressed as Mean ± S.D (n=3).

**Figure 7:**
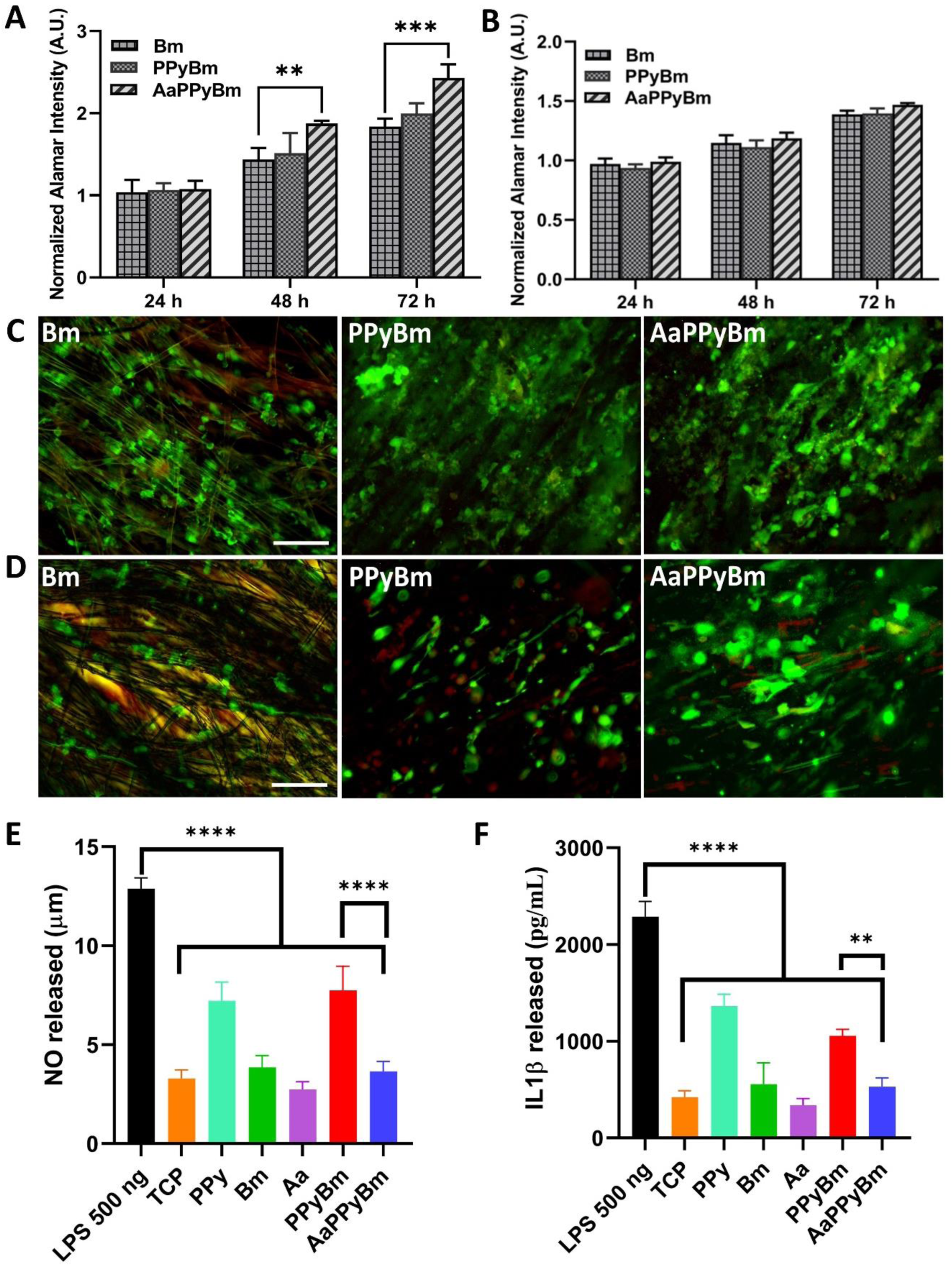
*In-vitro* biocompatibility results. Reduction of alamarBlue by the **(A)** primary porcine ADMSCs and **(B)** primary porcine DRG neurons cultured on different materials after 24, 48 and 72 h. Live-dead staining images showing live (green) and dead (red) **(C)** pADSCs and **(D)** pDRG neurons on different materials after 48 h, as indicated. *In-vitro* immune response results depicting **(E)** nitric oxide (NO) release after 12 h and **(F)** IL-β release after 24 h by Raw 264.7 macrophages, when treated with various materials. Data were Mean ± S.D, n=3.**≤0.01 and ***≤0.001 in **(A)** [Two-way ANOVA]; **≤0.01 and ****≤0.0001 in **(E)** & **(F)** [One-way ANOVA]; Scale bar = 200 μm (C & D).

Representative live/dead images illustrated almost no dead pADSCs on all the scaffolds, whereas a negligible number of dead pDRG cells can be seen in PPyBm and AaPPyBm but with relatively higher cell density than the pure Bm scaffolds. pADSCs and pDRG cells exhibited consistent adherence and uniform spreading along the fiber direction across all silk-based aligned scaffolds, with a notably enhanced cellular activity on AaPPyBm. The results corroborated the evidence of the highest cell viability of AaPPyBm as revealed by alamrBlue reduction assay. This is further supported by enhanced live cell density and spreading when tested these scaffolds with MG63 cell line [**Figure S11**].

*In vitro* immunocompatibility of various scaffolds was evaluated using murine macrophages RAW 264.7 cells stimulation with lipopolysacchariches (LPS) to predict the materials’ potential inflammatory response. The pro-inflammatory nitric oxide (NO) secretion was monitored after 12 h, while interleukin (IL-1β) secretion was recorded after 24 h using lipopolysaccharide (LPS) as positive control and non-stimulated tissue culture plate (TCP) as negative control [**Figure 7(E&F)**]. The results indicated AaPPyBm exhibited significantly lower pro-inflammatory NO and IL-1β secretion when compared to those with PPyBm and positive control (LPS). In fact, the pro-inflammatory NO and IL-1β modulators secretion by the immune cells on AaPPyBm is comparable to the negative control (TCP). However, PPy NPs coated silk scaffold without AaSF functionalization elicited higher pro-inflammatory response (but less than the positive control). The results show that functionalization of the PPyBm scaffolds with Aa did not contribute to the proinflammatory behavior. These *in vitro* biocompatibility evaluations substantiate the hypothesis that introducing Aa functionalization would improve cellular behavior on PPy:Silk-based scaffolds while minimizing immunomodulatory responses.

### 3.7 Neural differentiation of primary ADMSCs

The PPy:Silk based aligned conductive scaffolds displayed biologically relevant features such as in vitro biocompatibility, tuneable biodegradability along with decent physicochemical properties in terms of electrical conductivity and mechanical strength that matches with nerve tissue. This has placed these materials as smart biomaterial for electrically stimulated accelerated nerve regeneration. Hence, to assess the potential in nerve regeneration, neuronal like differentiation of primary pADSCs were carried out on these conductive scaffolds following the previous protocols^24, 110, 111^. We investigated to check whether the aligned morphology along with enhanced electroconductivity and bioactivity of the scaffolds after functionalization with AaSF, exert any favorable influence on neuronal like differentiation of primary pADSCs. To verify neuronal like differentiation of pADSCs on the various PPy:Silk scaffolds, cells were subjected to immunostaining with β (III) tubulin, a key early neuron specific marker for immature neurons and GFAP, a glial cell specific marker, following a 14 day culture period [**Figure 8(A&C)**]. Cells were also counterstained with phalloidin conjugated to rhodamine and hoechst for F-actin and nucleus, respectively. The F-actin dynamics (cytoskeletal organization) is pivotal in governing the motility of growth cones, making it a crucial factor in the processes of axon growth and branching^112^. Cytoskeleton rearrangement was revealed through rhodamine phalloidin staining where cells were aligned along the fiber direction, displaying extended protrusions rich in actin or filopodial outgrowth originating from the cell body. This phenomenon is instrumental in the formation of growth cones, facilitating neurite-like extensions. Prior studies have documented the capacity of a smaller fraction of pADSCs to undergo neural supporting Schwann cell or glial cell differentiation under the similar differentiation protocol, alongside functional neuronal commitment by the larger fraction^113^. Consistent with this observation, the current study reveals a substantial proportion of cells expressing the neuron-specific β (III) tubulin within microtubule assemblies, while a relatively lower number of pADSCs were positive for glial specific GFAP marker. All the staining images reveal that cells adhered and spread along the fiber direction following contact guidance phenomenon on all the aligned scaffolds. Quantitative assessment of the immunostaining results further shows that the higher numbers of cells differentiated towards neuronal lineage [expressing β (III) tubulin] and glial phenotype [expressing GFAP] on AaPPyBm, when compared to that on PPyBm and pure Bm scaffold [**Figure 8(B&D)**]. The enhanced neuronal or glial differentiation on AaPPyBm can be ascribed to the better cell-to-cell communication due to the excellent cell adhesion and enhanced electronic conductivity offered by AaSF functionalization. In fact, the nanoparticle features of PPy along with their conductive properties has a major role in cytoskeletal organization for neuronal differentiation, which is in agreement with previous findings^24, 114^. Nanostructured morphology present in biomaterial scaffolds can influence integrin clustering and the assembly of focal adhesions, which plays a crucial role in inducing changes in cellular morphology. This is also supported by elevated differentiation of pADSCs on PPyBm than the pure Bm scaffolds.

**Figure 8:**
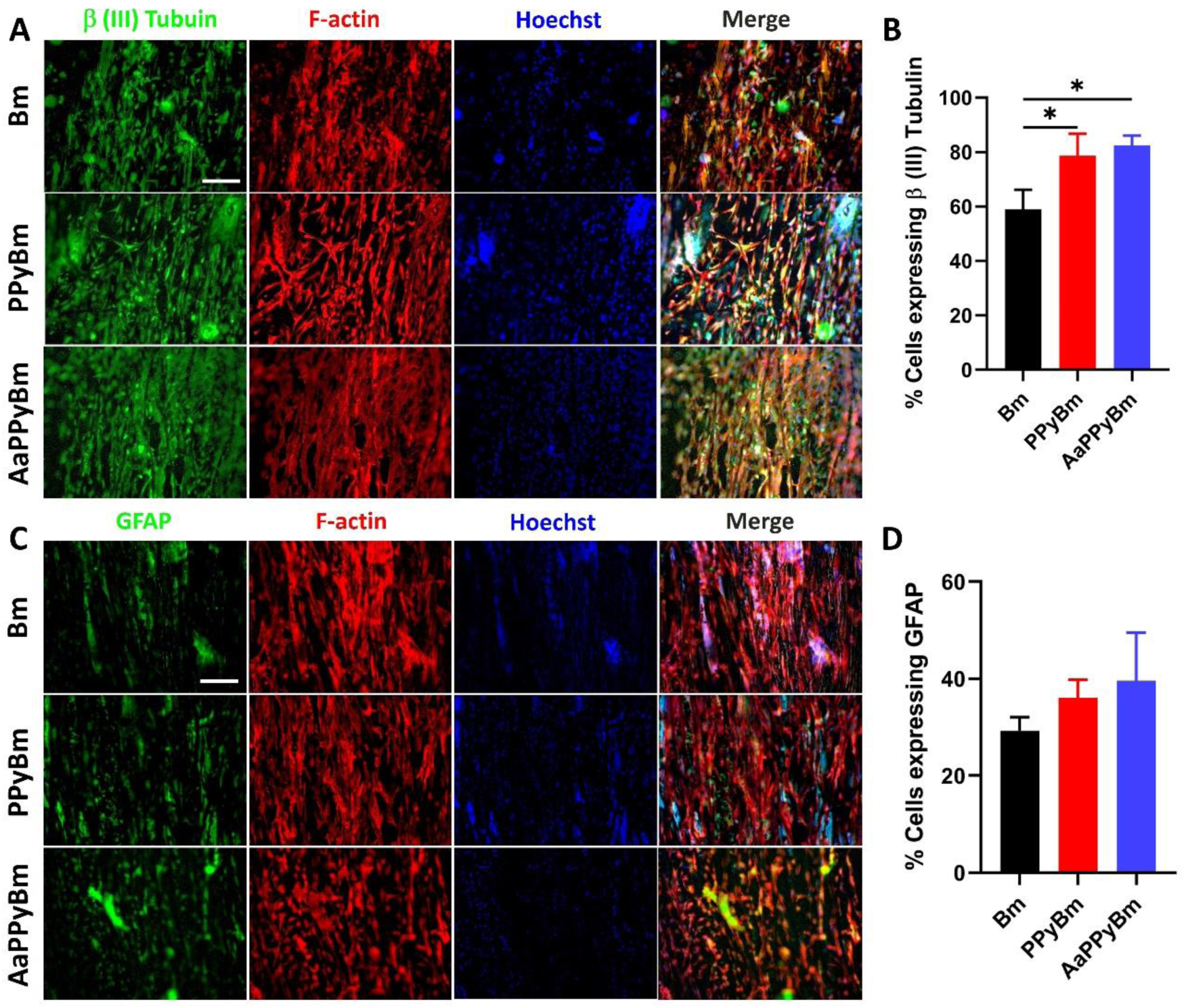
Immunostaining results revealing the ability of various silk-based aligned scaffolds to promote neuronal like differentiation of primary porcine pADSCs. Representative fluorescence micrographs of cells immunostained by **(A)** neuronal marker β (III) tubulin antibody (green), and **(C)** glial cell specific GFAP antibody (green) counterstained by rhodamine-phalloidin for F-actin filaments (red) and hoechst for nucleus (blue) after 14 days of culture in differentiating medium. Quantitative analysis showing % cells expressing **(B)** β (III) tubulin neuronal marker and **(D)** glial cell marker GFAP. *corresponds to statistical significance at p≤0.05 in **(B)** [One-way ANOVA]; Scale bar = 200 μm.

### 3.8 Electrically stimulated axonal growth of primary pDRGs

Both PPyBm and AaPPyBm scaffolds showed enhanced neuronal-like differentiation of pADSCs as indicated by immunostaining results owing to their good intrinsic electrical conductivity, as discussed earlier. However it is noteworthy that AaPPyBm shows longer neurite projections compared to PPyBm. This observation can be correlated with PPy’s unique ability to integrate large anionic biomolecules, such as hyaluronic acid (HA) and chondroitin sulfate (CS)^115^, or amino acids like lysine^71^ and glutamate^72^, and even the RGD peptide^73^, as dopants within its network, ultimately enhancing its conductive behaviorMoreover, electroneutrality coupling and electron-hopping mechanisms in PPy are affected when it encounters different anions, enhancing its conductivity and electrochemical behaviour^116^. Therefore, in this study, AaSF functionalization may also alter the charge density within PPy:Silk scaffold, facilitating the Faradaic charge injection/transfer process at electrode-tissue interface as evidenced by the low charge transfer resistance of AaPPyBm [**Figure S15 &Table 3**]. Hence, it can be summarized that the improved bioactivity from RGD tripeptide and altered surface charge chemistry due to AaSF functionalization resulted in enhanced nerve growth on AaPPyBm. The interaction of anionic AaSF with the PPy chain to make it more redox active with electron delocalization is also confirmed by FT-IR, Raman, and Zeta potential analysis. However, the exact effect of scaffold conductivity along with their improved electrochemical performance is anticipated to be more pronounced in electrically stimulated neurite growth^117^. To validate our hypothesis that the scaffolds exhibiting elevated CSC, current/charge injection capacity, and reduced electrochemical charge-transfer resistance (R_ct_), can accelerate axonal growth at lower stimulation voltage, the neurite growth characteristics of primary pDRG sensory neurons seeded on various conductive scaffolds subjected to pulsed ES, were evaluated. Pulsed ES (50 Hz,1 ms) with varying amplitude (50, 100, 200, and 300 mV cm^-1^) was administrated to the cells through the various conductive scaffolds for 2 h daily over 3 consecutive days and the representative waveform of pulsed ES is shown in **Figure S14**. The cell-seeded scaffolds were stained with β (III) Tubulin neuronal marker after 7 days of culture to visualize the expression of neuronal characteristics. Representative fluorescent images of non-stimulated and stimulated β (III) Tubulin stained pDRG neurons on various conductive scaffolds are shown in **Figure 9 (A)**.

**Figure 9:**
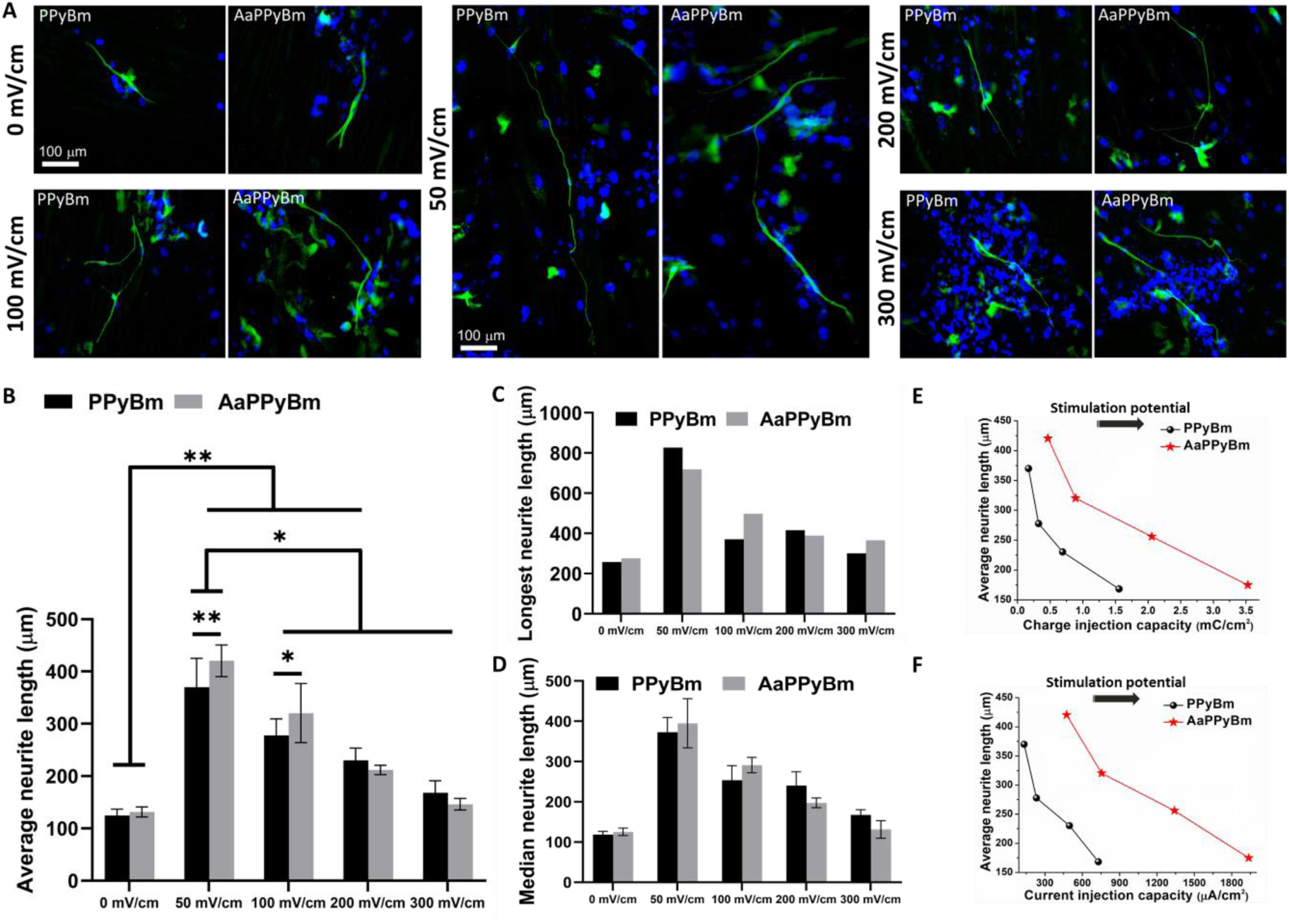
Electrical stimulation (ES) results. **(A)** β (III) tubulin immunostaining (green) of dorsal root ganglions (pDRGs) neurons, counterstained with hoechst for nucleus (blue), grown on the PPy:Silk scaffolds after 7 days under pulsed ES of frequency 50 Hz, pulse width 1 ms with different amplitudes from 50-300 mV cm^-1^ for 2 h/day (3 days), as indicated (Scale bar = 100 μm). Quantitative assessment of ES results showing **(B)** average neurite length, **(C)** longest neurite length and **(D)** median neurite length. Variation of neurite outgrowth rate with respect to **(E)** injected charge (charge injection capacity) & **(F)** injected current (current injection capacity) at different stimulation potentials, depicting effect of material’s intrinsic electrochemical properties on neurite growth. * and ** indicate statistical significance at p≤0.05 and p≤0.01 [Two-way ANOVA].

The images reveal neurite projection or elongation following fiber alignment in all scaffolds, with ES-treated axons being notably longer than those without ES. In addition to the sensory neurons, DRG culture contains various cell types, including neural supporting cells and fibroblasts, as indicated by nucleus staining (blue) in the β (III) tubulin-stained images. The presence of non-neuronal cells is shown in dissociated DRG neuronal cultures [**Figure S13]**. The β (III) tubulin-stained images were analyzed using ImageJ software for the quantitative evaluation of the neurite growth axonal length measured as the linear distance between the cell junction and neurite tip. Axonal length data of at least twice the diameter of the cell body were chosen for analysis, with around N=80-150 neurites analyzed per sample.

The results show that ES treated cells demonstrate significantly longer neurite outgrowth ( measured as average, median, and longest neurite length) compared to unstimulated cells [**Figure 9(B-D)**]. Interestingly, the neurite projections are the longest at a low stimulation potential of 50 mV cm^-1^, in contrast to those at relatively higher stimulation potentials [**Figure 9(B-D)**]. The neurite projections are the shortest at the stimulation potential of 300 mVcm^-1^, and at 500 mV cm^-1^, most cells appear dead (Results not shown).

Two-way ANOVA analysis demonstrates that the average neurite length at 50 mV cm^-1^ (on both PPyBm and AaPPyBm) is statistically significant compared to other stimulation potentials (p≤0.05). The neurite outgrowth extends up to 830 μm at 50 mV cm^-1^, compared to the longest neurite lengths of 300-500 μm at 100-300 mV ^-1^cm or 275 μm at 0 mV cm^-1^ [**Figure 9(D)**]. This observation can be linked to the intrinsic electroactivity of the PPy:Silk scaffolds. Specifically, the observed trend can be correlated with the charge/current injection limit of the scaffolds during ES and their CSC. A study by Abidian *et al.* supports the current observation, showing enhanced neurite outgrowth of DRG explants on PPy and PEDOT nanotubes with CSC of 184 and 392 mC cm^-2^.^109^ The PPy:Silk based scaffolds demonstrated excellent CSC due to their high electrical conductivity (6.23-9.18 mS cm^-1^) [**Table 2**] and porous morphology, consistent with previous studies.^7, 118^ This implies that the scaffolds can accommodate a sufficient quantity of charge carriers to interact with cell membranes during ES., Additionally, they display enhanced charge injection capacity ranging from 0.16 mC cm^-2^ at 50 mV to 3.52 mC cm^-2^ at 300 mV with 1 ms pulse width, indicating improved stability revealed by chronoamperometric analysis in NBM [**Figure 5(E)**]. ES was also conducted under identical conditions, maintaining the same pulse width and amplitude, so the injected charge/current was assumed to be equivalent during the process. The injected charge limits for both PPyBm and AaPPyBm are well within the safe limit for human or animal tissues, particularly at lower stimulation potentials at 50 and 100 mV cm^-1^ and are sufficient to stimulate neurons. Due to their excellent conductivity, the scaffolds were also able to deliver 100-700 μA of current at 50 and 100 mV cm^-1^ [**Figure 5 (F)**]. This explanation aligns with a prior study, where a charge density of 0.05-0.1 mC cm^-2^ or 5 μA current was successfully applied to evoke a retinal ganglion spike.^5^ However, when the ES exceeded 100 mV cm^-1^, the neurite outgrowth rate significantly decreased, owing to the increase in the injected charge and current density [**Figure 9(E&F)**]. A similar observation was reported by Rajnicek et. al., where they showed varied electrochemical behaviour of different conducting scaffolds, viz., PEDOT:PSS, IrOx and mixed oxide (Ir-Ti)Ox, during ES, significantly influenced neurite growth^119^. The study showed sufficient neuronal growth at a low stimulation potential of 50 mV mm^-1^ without material delamination, while at higher potentials of 100-150 mV mm^-1^, reduced neurite growth was linked with the disintegration of material adhesion due to overoxidation. It is worth noting that very high ES potential, particularly with hybrid conductive scaffolds, can lead to elevated charge carrier transfer and electrochemical interactions resulting in over-oxidation or reduction at the scaffold-tissue interface. In the present study, the notion is that at low stimulation potential, charge-transfer occurs mainly at the surface of the electrodes through faradaic processes, while this process can exceed the interior of material resulting in material degradation at higher stimulation due to increased charge injection. Higher charge injection may also result in oxidation/reduction of the cell culture medium, forming oxygen free radical, and gases like H_2_ and O_2_.Consequently, this can cause several harmful effects in neuronal dysfunction or damage through excessive activation of the excitatory neurotransmitter (excitotoxicity), excessive ion flow leading to ion imbalance across the cell membranes, oxidative stress, and neuronal hyperactivity. Thus, the neuronal projections decrease with the increase in the stimulation potential, confirmingthe effective functionality of these hybrid electroactive scaffolds at lower stimulation potentials.

Furthermore, neurite extension on AaPPyBm is significantly greater than on PPyBm at ES of 50 and 100 mV cm^-1^. AaPPyBm, which possess better CSC, current/charge injection capacity and lower R_ct_ and superior electrical conductivity (∼9.18 mS cm^-1^), showed better efficacy in ES mediated neurite growth. Previous studies also showed that scaffolds with higher electrical conductivity and charge carrier density elicited enhanced neural differentiation of PC12 cells and accelerated neurite outgrowth under ES.^22^.Rajnicek et. al. reported that an indirect electrical potential or dipole is induced within the conductive material during ES through a phenomenon called “bipolar electrochemistry”.^119^ This induced electric potential originates at the scaffold-media interface, whose nature depends on material’s intrinsic surface charge and plays a crucial role in nerve growth during ES. On the other hand, due to higher charge injection along with higher charge transfer efficiencylead to reduced neurite outgrowth on AaPPyBm than PPyBm. These results suggest that such hybrid electroactive scaffolds with enhanced conductivity and electrochemical properties can effectively promote neural regeneration at relatively lower stimulation potential, which is desirable for a safe nerve stimulation protocol. However, there is further scope in assessing the molecular mechanisms and *in vivo* functionality of these NGCs developed here, which is the basis for further investigation. Moreover, combining these scaffolds with drug delivery systems or advanced fabrication techniques, such as 3D printing, could open new avenues for precision nerve repair.

## 4. Conclusions

The current study shows the fabrication of a biohybrid electroactive aligned PPy:Silk based microfibrous scaffold, functionalized with unique Indian no-mulberry AaSF containing the cell-affinitive RGD tripeptide, capable of inducing a functional neuronal stimulation at low voltages. The study demonstrated, for the first time, the positive impact of highly bioactive AaSF on nerve growth. Furthermore, detailed investigation uncovered a secondary role for anionic AaSF as a dopant for PPy, which led to enhanced electrical conductivity, charge density, and charge-transfer efficiency of the PPy:Silk scaffold. Surface morphological analysis revealed increased interconnectivity among the PPy NPs after AaSF functionalization, which also consequently made the scaffold mechanically stronger maintaining adequate flexibility as shown by tensile studies. Notably, our optimized protocol for in-situ polymerization of PPy NPs produced highly conductive PPy:Silk scaffold with conductivity of 6.23 mS cm^-1^, significantly surpassing the results of most previous studies. This was further enhanced upto ∼9.18 mS cm^-1^ after AaSF modification of the scaffold, indicating its potential role as an anionic dopant. The biohybrid scaffolds displayed comparable or higher charge injection capacity, even at low potentials (e.g., 0.46 mCcm^-2^ at 50 mV), compared to conventional bioelectrodes. This indicates their potential to deliver therapeutic electrical stimulation at lower electrical potentials, without causing electrode degradation or tissue damage. This is again corroborated with the extended neurite outgrowth of DRG neurons upto 830 μm, subjected under pulsed ES of amplitude 50 mV cm^-1^ through these biohybrid scaffolds. However, the rate of neurite outgrowth decreases as the injected current/charge density increases, along with a corresponding rise in stimulation voltage. Thus, the investigation of electrically stimulated neurite growth characteristics in relation to the scaffolds’ CSC, charge/current injection capacity, and charge-transfer efficiency, highlights the importance of an optimum injected charge/current density at low stimulation voltages for evoking a functional action potential for membrane depolarization. The biohybrid scaffolds also demonstrated enhanced neuronal or glial differentiation owing to the better cell-to-cell interaction due to the excellent cell adhesion and enhanced electronic conductivity offered by AaSF functionalization. To conclude, the study demonstrates a strategy to predict the suitable ES parameters for electrically excitable tissues or cells while highlighting the importance of the scaffold’s intrinsic electronic or electrochemical properties to electrically stimulated neurite growth. Tuning such material property can be extremely useful using biologically active molecules such as AaSF, which enhances the bioactivity of the PPy:Silk scaffold in addition to improving their electronic and electrochemical properties. Thus, the fabricated biohybrid scaffolds hold great potential for future implications as smart nerve guidance channels (NGCs).

## Acknowledgements

RB and BBM gratefully acknowledge the financial support provided by the Department of Science and Technology (DST), Government of India (Grant ID: DST/INSPIRE/04/2018/000402), and partial funding from IIT Guwahati through the institutional post-doctoral fellowship program (Grant ID: IITG/R&D/IPDF/2017-2018/BSBE01). JU acknowledges the Science and Engineering Research Board (SERB), Government of India (Grant No. ECR/2017/000628), for funding the electrochemical measurements reported in this work. We also extend our sincere thanks to the Sophisticated Analytical Instrument Centre (SAIC) at IIT Guwahati and the SAIC of IASST for their invaluable assistance with the physicochemical characterization of the materials.

