## Supporting information for "Non-mulberry silk fibroin functionalization enhances charge-transfer efficiency in aligned polypyrrole-silk composites for electrically stimulated neurite outgrowth"

### ELECTRONIC SUPPLEMENTARY INFROMATION

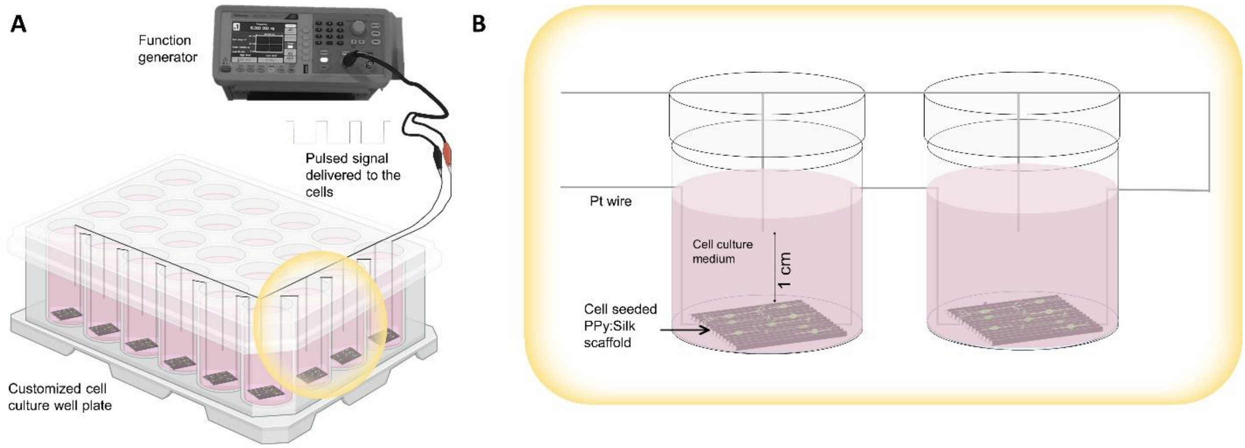


**Figure S1**: Schematic illustration of (A) bespoke electrical stimulation (ES) set up and (B) interconnected cell culture wells with scaffolds through Pt wires for ES.

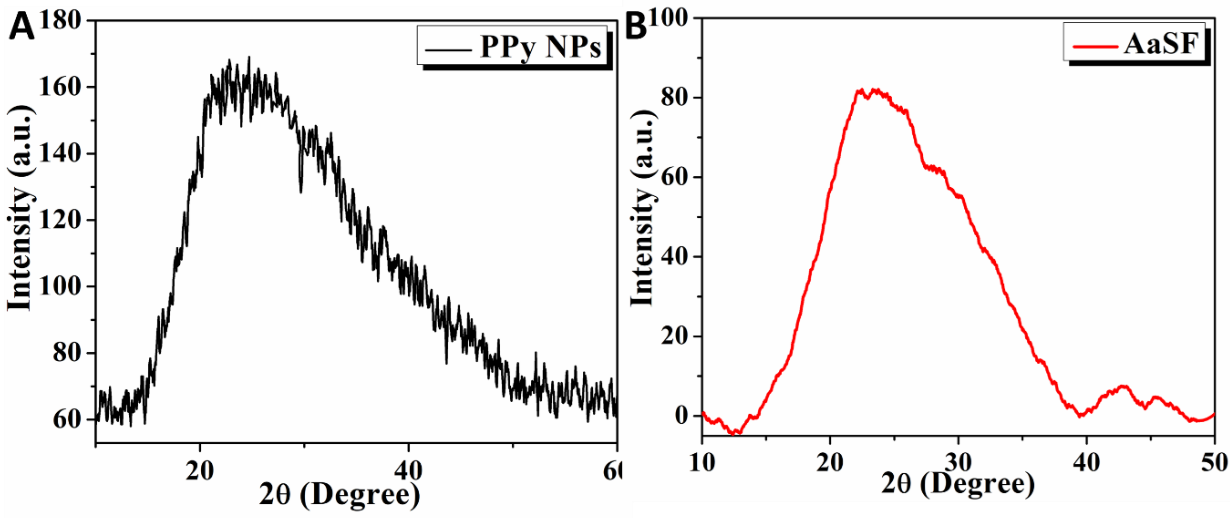


**Figure S2**: X-ray diffraction pattern of (A) PPy NPs and (B) freeze-dried *Antheraea assamensis* silk fibroin (AaSF) powders.

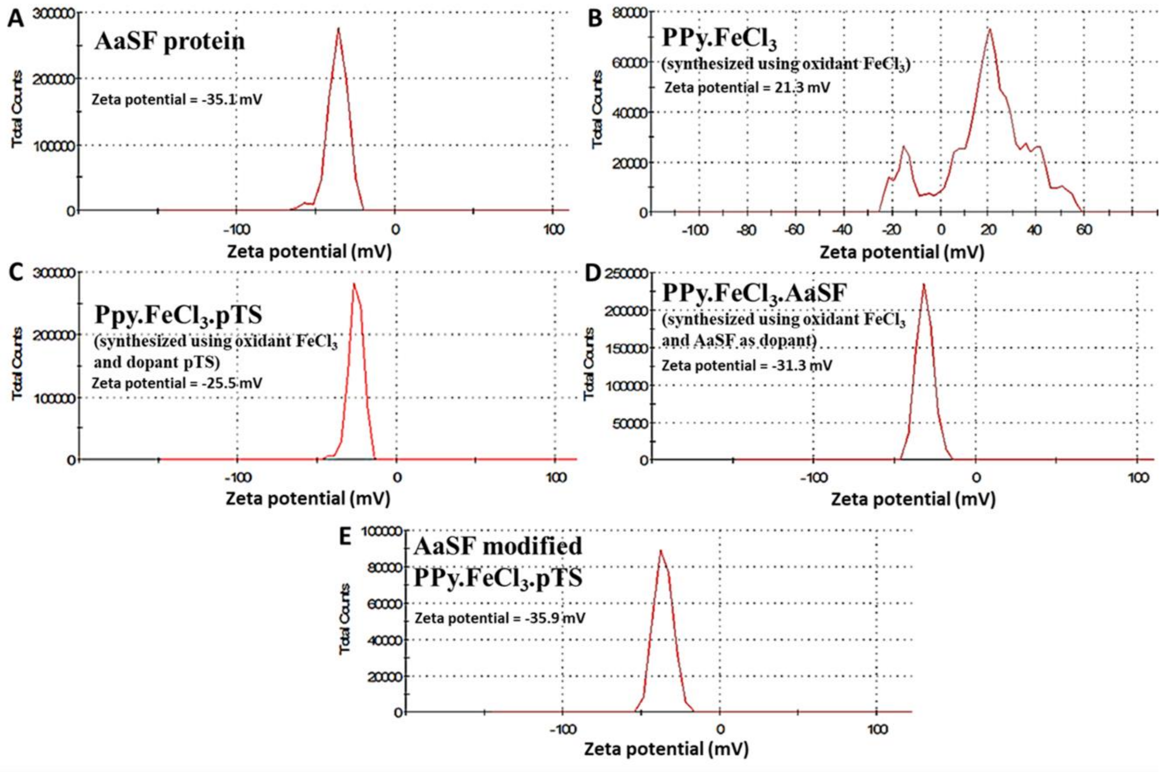


**Figure S3**: Zeta potential distribution of (A) AaSF protein, (B) PPy NPs synthesized using FeCl3 as oxidant (no dopant), (C) PPy NPs synthesized using FeCl3 and pTS as oxidant and dopant, respectively, (D) PPyNPs synthesized using $FeCl_3$ and AaSF as oxidant and dopant, respectively, and (D) AaSF functionalized PPy NPs synthesized using $FeC_{l3}$ and pTS as oxidant and dopant, respectively. All measurements were performed in milliQ water (pH=6.9-7.2).

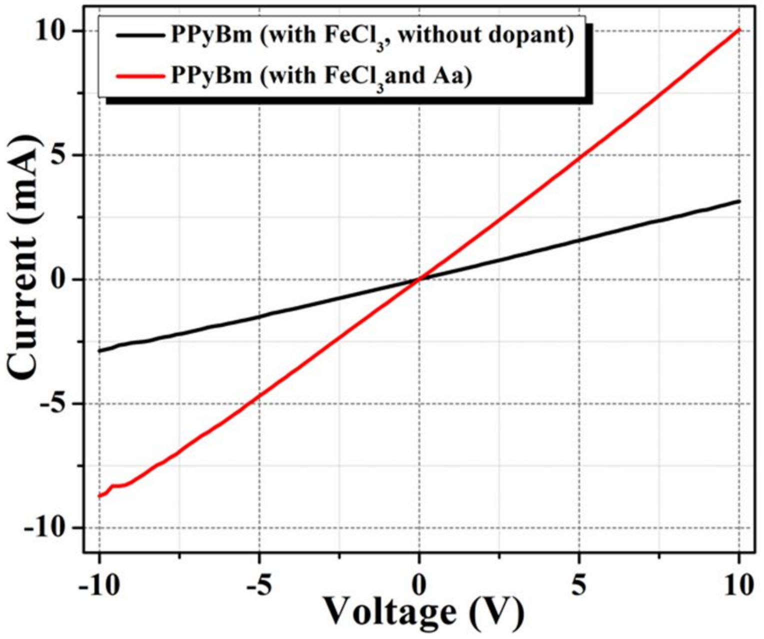


**Figure S4**: Room temperature (300 K) current-voltage (I-V) characteristics of two variants of PPyBm; PyBm fabricated with $FeCl_3$ as oxidant and without using any dopant (black); PPyBm fabricated with FeCl3 and AaSF as oxidant and dopant, respectively, (red).

**Table S1**. Some attempts on use of silk fibroin and poly(pyrrole)-based composites for developing electroconductive biomaterials (with impetus towards neural tissue engineering).

| **Material** | **Method** | **Electrical/ Electrochemical properties** | **Key Findings** | **Reference** |
|---|---|---|---|---|
| PPy coated silk fabrics | In-situ chemical polymerization | - | • The conductivity of PPy-coated silk fibers decayed upon exposure to environmental conditions.<br>• 10-20% conductivity decreases over a period of 2 years.<br>• Cellular performance not validated<br>• Composite yarns remained conductive for 2-year period | Boschi et al. 2020[1] |
| Silk mesh coated with PPy | Electrospinning & chemical polymerization | - | • No additional bioactive coating is done.<br>• PPy coated meshes allow anion storage during redox reactions, making them useful for biosensing and drug delivery applications.<br>• Cytocompatibility assessments using fibroblasts and mesenchymal stem cells done. No neural activity using neuronal cells studied. | Aznar-Cervantes et al 2012[2] |
| 3D silk foam poly(pyrrole-co-(2-hydroxy-5-sulfonic aniline)) scaffold | Salt-leaching (porogen method) to make porous foams & chemical polymerization | $6.1 \times 10^{-4}$ S | • Scaffolds facilitate electrical stimulation of mesenchymal stem cells, promoting their differentiation into osteogenic outcomes.<br>• Foams were highly conductive (~0.6 mS/cm) matching the conductivity of mammalian tissues (≥0.1 mS/cm). | Hardy et al 2015[3] |

| | | | | |
|---|---|---|---|---|
| | | | • Tested for osteogenic differentiation only | |
| PPy modified gelatin-silk fibroin scaffold | Solvent casting films & in-situ chemical polymerization | $R_{ct}$ ~ 279-1386 $\Omega.cm^2$ | • The conductivity of scaffolds improves as the PPy content increases.<br>• Improved charge-transfer behavior<br>• Cellular performance not validated | Kulkarni et al 2020[4] |
| Acid modified silk-poly(pyrrole) scaffolds | Chemical polymerization | ~1 S $cm^{-1}$ | • Enhanced expression and polarization of connexin 43 (Cx43), a key regulator of electrical coupling between cells.<br>• The substrates were biocompatible and stable, supporting high viability of hPSC-derived cardiomyocytes throughout a 21-day culture period. | Tsui et al 2018[5] |
| PPy-silk fibroin scaffold | 3D bioprinting and electrospinning | $1 \times 10^{-3}$ S $cm^{-1}$ | • The polypyrrole/silk fibroin scaffold demonstrated no cytotoxicity.<br>• The scaffolds exhibited good compatibility with L929 cells and have the potential to enhance Schwann cell adhesion, differentiation, and proliferation. | Zhao et al 2018[6] |
| PPy-silk composite | Polymerization of PPy over functionalized silk film | ~1 S $cm^{-1}$ | • The acid-modified scaffold exhibited increased stability against biodegradation when exposed to protease XIV and collagenase.<br>• The composites showed stable electrochemical performance. | Romero et al 2013[7] |

| | | | | |
|---|---|---|---|---|
| PPy-silk fibroin composite | 3D bioprinting and electrospinning | 0.11446 ± 0.00145 mS $mm^{-1}$ | • To enhance cell attachment 0.01% PLL was used for coating<br>• Electrical stimulation enhanced neurite outgrowth and facilitated neural network formation.<br>• The MAPK signaling pathway was activated by electrical stimulation. | Zhao et al 2020[8] |

**Table S2:** FT-IR peak positions of various silk-based scaffolds showing their position shifting with description.[5, 7, 9-16]

| Peak position ($cm^{-1}$) |
|---|

| Chemical bond, group etc. | Bm | Aa | PPy | PPyBm | AaPPyBm |
|---|---|---|---|---|---|
| **C-H out of plane bending** | 1015-1057 (weak to medium) | 1021-1082 (weak to medium) | 1037-1114 (strong to weak); Overlapped with C-H bending of pyrrole ring and S=O stretching in aryl sulfonate salt suggesting the doping level in PPy; Distinct band at 966 cm-1 refers to C-H/C-C bending unaffected by dopants | 1032-1114 (strong to weak) | 1035-1116 (strong to weak); Relatively weak band at 962 $cm^{-1}$ indicates increased oxidation after functionalization with Aa |
| **Amide III/C-N stretching in PPy** | 1233 (strong) & 1333 (medium); mixed C-N stretching and N-H bending of amide III | 1214 (strong), 1308-1336 (weak) | 1163 (strong)-C-N stretching in pyrrole ring; 1310 (strong)-C-N stretching in plane deformation; 1359 (weak)- C-$N^+$ stretch with C-C vibration | 1150 & 1296 (strong); C-$N^+$ stretch at 1362 $cm^{-1}$ is weak but distinct; Overlapped with C-N stretching & N-H bending of amide III | 1158-1300 (strong & broad); Distinct C-$N^+$ stretch at 1362 $cm^{-1}$; Contribution from amide III |
| **C-C stretching & overlapping vibrations** | 1385-1407 (weak) & 1448 (medium) | 1383-1412 (weak) & 1452-1463(broad & medium) | 1414 (weak) & 1467 (medium, broad)-PPy ring breathing with contributions from C-C/C=C and C-N | Multiplets at 1414-1472 (weak to strong)-PPy ring vibration | Distinct multiplets at 1417-1477 (weak to strong); Weak bands at 1382-1400 refers to SF contribution |
| **Amide II/N-H bending in amine & overlapping vibrations** | 1509-1541 (broad, strong to medium) due to amide II | 1511-1539 (broad, medium to strong) due to amide II | 1548 (strong)-C=C/C-C stretching along with N-H bending in secondary amine of PPy; 1598 (weak)-C=$N^+$ vibration of half oxidized pyrrole | 1502-1538 (strong); Overlapping of amide II and PPy ring vibration | 1508-1560 (broad & strong); Overlapping of amide II and PPy ring vibration |

| | | | | | |
|---|---|---|---|---|---|
| **Amide I** | 1620 (strong)-amide I of β-sheet of SF; 1654 (shoulder peak)-amide I of α-helix/random coil of SF; C=O stretching in amide I with minor contribution from N-H in pane bending, the out-of-phase C≡N stretching vibrations and the C−C≡N deformation | 1652 (strong)-amide I of α-helix/random coil | - | 1622 (strong) & 1650/ 1682/1692 (weak); Due to conformational changes in protein secondary structure due to interaction of PPy with polypeptide chain of SF; Weak signals at 1690-1720 $cm^{-1}$ due to oxidized PPy | 1617 (strong) & 1663 (strong)-contributions from amide I in PPyBm and Aa; Distinct band at 1698 $cm^{-1}$ due to C=O stretching due to -COOH group (of Aa) conjugated with secondary amine of PPy; Overlapped with other bands at 1700-1730 $cm^{-1}$ due to overoxidized PPy |
| **C-O stretching** | - | - | - | 2319-2348 (strong) | 2322-2355 (strong) |
| **C-H stretching** | 2930-3071 (weak) | 2850-3066 (strong to weak) | 3043 (broad. weak) | 2835-2954 (weak) | 2838-2972 (weak to medium) |
| **N-H stretching/O-H stretching** | 3278 (strong, broad) | 3292 (strong, broad) | 3190 (broad, weak) due to N-H stretching | 3275 (weak, broad); Overlapping of N-H and O-H stretching | 3428 (medium, broad); Overlapping of N-H and O-H stretching |

**Table S3:** The electrochemical behavior of the scaffolds depicting their redox activity both in neurobasal media (NBM) is summarized with their redox potentials and peak currents (anodic & cathodic).

| NBM | | | | | | | | | | | | |
|---|---|---|---|---|---|---|---|---|---|---|---|---|
| **Scan rate ($mV.s^{-1}$)** | Anodic peak potential ($E_{pa}$) (mV) | | | Cathodic peak potential ($E_{pc}$) (mV) | | | Anodic peak current ($I_{pa}$) ($mA.cm^{-2}$) | | | Cathodic peak current ($I_{pc}$) ($mA.cm^{-2}$) | | |
| | PPyBm | AaPPyBm | | PPyBm | AaPPyBm | | PPyBm | AaPPyBm | | PPyBm | AaPPyBm | |
| **10** | 238 | 83 | 232 | -460 | -327 | -400 | 1.05 | 0.89 | 1.20 | -1.49 | -1.52 | -1.55 |
| **50** | 172 | - | 168 | -436 | -357 | -447 | 0.97 | - | 1.42 | -1.78 | -2.00 | -2.26 |
| **100** | 232 | - | 264 | -463 | -508 | -601 | 1.35 | - | 1.68 | -1.97 | -2.43 | -2.51 |
| **150** | 249 | - | 297 | -468 | -462 | -588 | 1.50 | - | 2.09 | -2.01 | -2.78 | -2.81 |
| **200** | 253 | - | 298 | -481 | -457 | -598 | 1.55 | - | 2.26 | -2.14 | -2.95 | -3.17 |

**Table S4:** The electrochemical behaviour of the scaffolds depicting their redox activity both in PBS is summarized with their redox potentials and peak currents (anodic & cathodic).

| PBS | | | | | | | | | | | | |
|---|---|---|---|---|---|---|---|---|---|---|---|---|
| **Scan rate ($mV.s^{-1}$)** | Anodic peak potential ($E_{pa}$) (mV) | | | Cathodic peak potential ($E_{pc}$) (mV) | | | Anodic peak current ($I_{pa}$) ($mA.cm^{-2}$) | | | Cathodic peak current ($I_{pc}$) ($mA.cm^{-2}$) | | |
| | PPyBm | AaPPyBm | | PPyBm | AaPPyBm | | PPyBm | AaPPyBm | | PPyBm | AaPPyBm | |
| **10** | -24 | 48 | - | -233 | -304 | -502 | 0.21 | 3.23 | - | -0.64 | -3.21 | -3.80 |
| **50** | -5 | 100 | - | -276 | -347 | -504 | 0.30 | 3.72 | - | -0.85 | -4.02 | -4.13 |
| **100** | 19 | 137 | - | -286 | -384 | -564 | 0.39 | 4.23 | - | -1.01 | -4.80 | -4.64 |
| **150** | 68 | 201 | - | -347 | -470 | -575 | 0.54 | 5.03 | - | -1.28 | -6.21 | -5.82 |
| **200** | 114 | 206 | - | -384 | -509 | -615 | 0.72 | 5.33 | - | -1.64 | -7.13 | -6.86 |

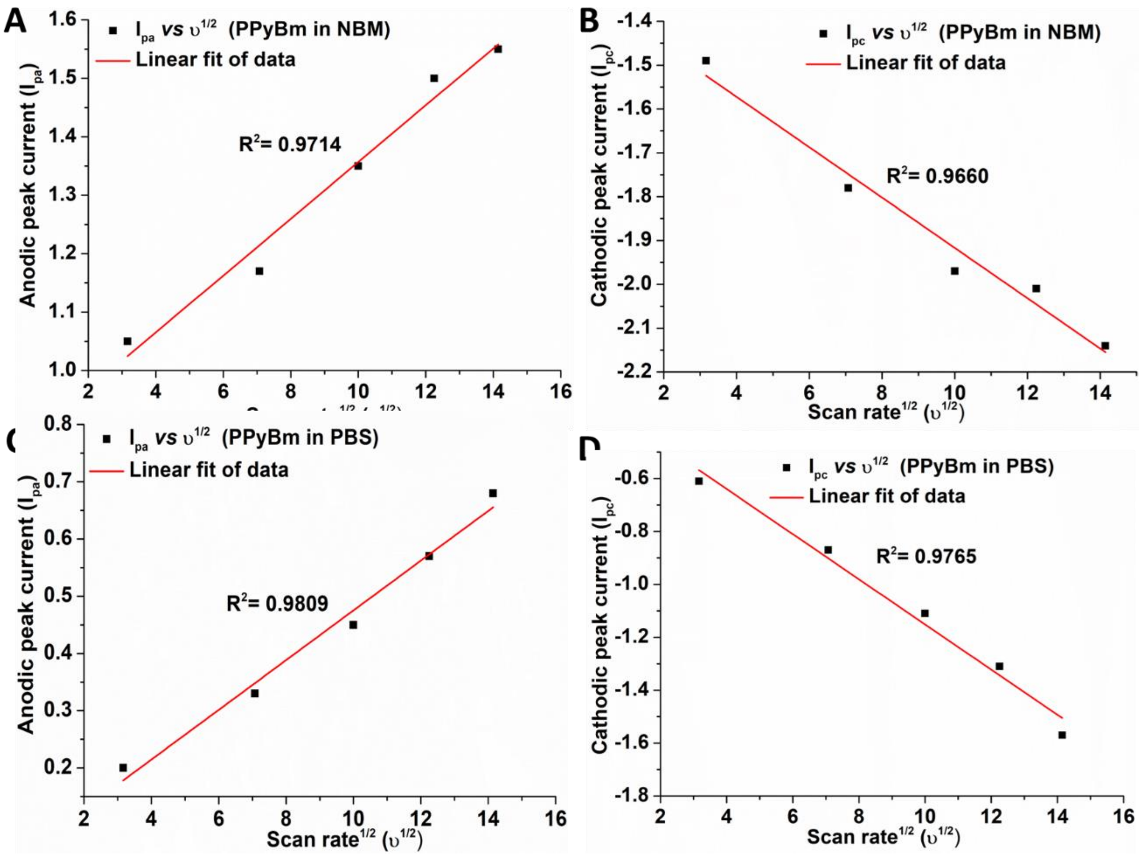


**Figure S5**: CV data fitting of PPyBm; The data depicts increase in the redox peak potentials for each scaffold in both the electrolytes with increasing scan rates (ʋ), while the redox peak currents vary linearly with $\upsilon^{1/2}$.

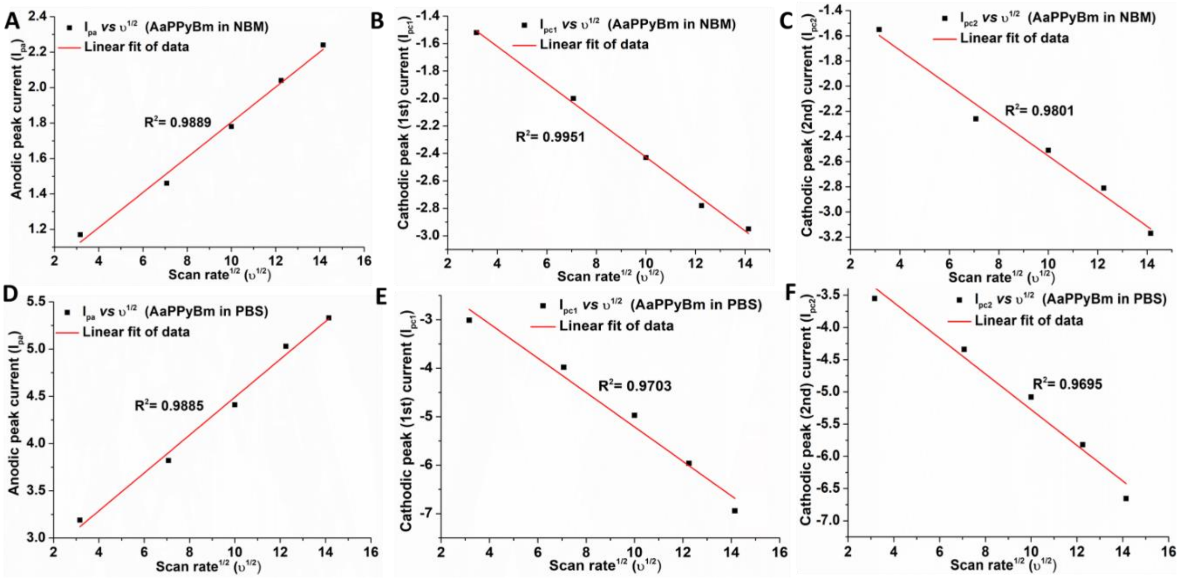


**Figure S6**: CV data fitting of AaPPyBm; The data depicts increase in the redox peak potentials for each scaffold in both the electrolytes with increasing scan rates (υ), while the redox peak currents vary linearly with $\upsilon^{1/2}$.

**Table S4:** Inorganic salt composition in Neurobasal medium (NBM)**.

| Inorganic Salts | Concentration (mM) |
|---|---|
| Calcium Chloride ($CaCl_2$) (anhyd.) | 1.8018018 |
| Ferric Nitrate ($Fe(NO_3)_3.9H_2O$) | 2.4752476E-4 |
| Magnesium Chloride ($MgCl_2$) | 0.8136842 |
| Potassium Chloride (KCl) | 5.3333335 |
| Sodium Bicarbonate ($NaHCO_3$) | 26.190475 |
| Sodium Chloride (NaCl) | 68.965515 |
| Sodium Phosphate monobasic ($NaH_2PO_4$-$H_2O$) | 0.9057971 |
| Zinc sulfate ($ZnSO_4$-$7H_2O$) | 6.736111E-4 |

****** Thermo Fisher, Catalogue number: 10888022

**Table S5:** Inorganic salt composition in 10 X PBS.

| Inorganic Salts | Concentration (mM) |
|---|---|
| Sodium Chloride (NaCl) | 8 g |
| Potassium Chloride (KCl) | 0.2 g |
| Sodium phosphate dibasic dihydrate ($Na_2HPO_4$) | 1.44 g |
| Monopotassium phosphate ($KH_2PO_4$) | 0.24 g |

**Table S6:** Hydrated ionic radius, ionic mobility and molar ionic conductivity of the electrolyte ions in DMEM and neurobasal media at 298K.[17-19]

| Ion | Radius of hydrated sphere (Å) | Ionic Mobility ($10^{-8}$ $m^2$ $s^{-1}$ $V^{-1}$) | Molar ionic conductivity (mS $m^2$ $mol^{-1}$) | NBM/PBS |
|---|---|---|---|---|
| $Na^+$ | 3.58 | 5.19 | 5.010 | Both |
| $K^+$ | 3.31 | 7.62 | 7.350 | Both |
| $Ca^{2+}$ | 4.12 | 6.17 | 11.90 | NBM |
| $Zn^{2+}$ | 4.3 | 5.47 | 10.56 | NBM |
| $Mg^{2+}$ | 4.28 | - | 10.60 | NBM |
| $Cl^-$ | 3.3 | 7.91 | 7.635 | Both |
| $CO_3^{2-}$ | 3.94 | 7.46 | 13.86 | NBM |
| $NO^{3-}$ | 3.35 | 7.40 | 7.146 | NBM |
| $SO_4^{2-}$ | 3.79 | 8.29 | 16.00 | NBM |
| $Fe^{3+}$ | 4.57 | NA | - | NBM |
| $PO_4^{3-}$ | 2.38 | - | - | Both |

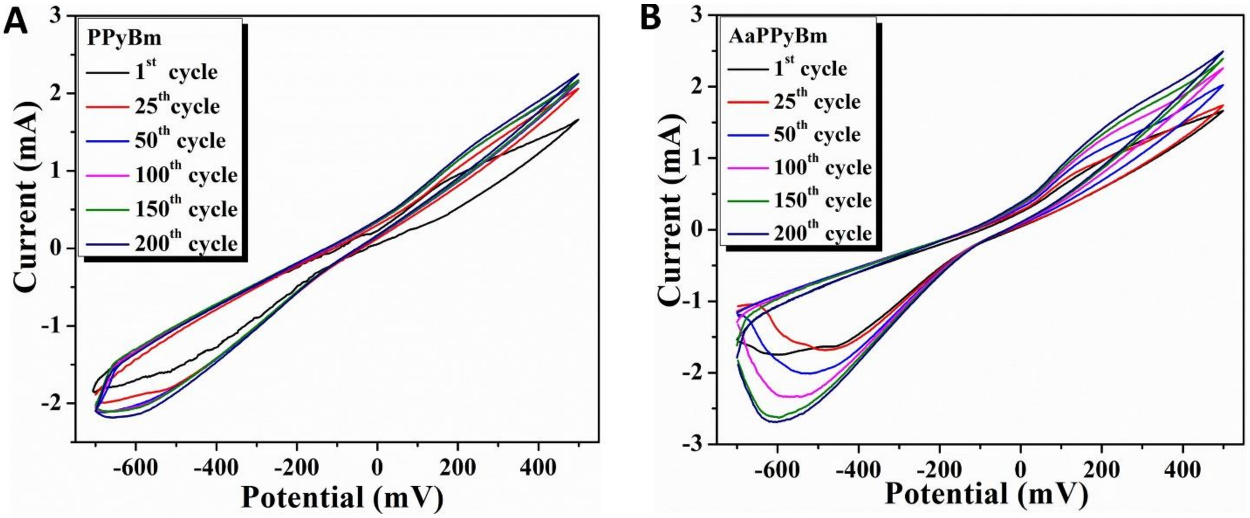


**Figure S7**: Cyclic stability study of **A.** PPyBm and **B.** AaPPyBm scaffolds up to 200$^{th}$ cycles.

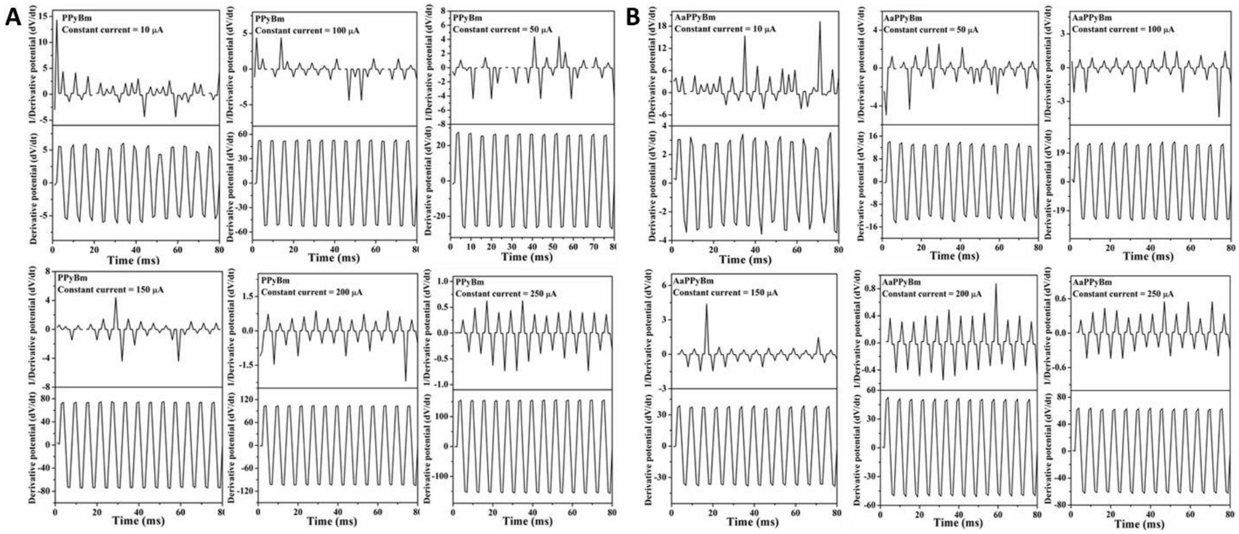


**Figure S8**: Derivative plots of the multiple chronopotentiometry responses and plots of their reciprocals at various current levels for (A) PPyBm and (B) AaPPyBm.

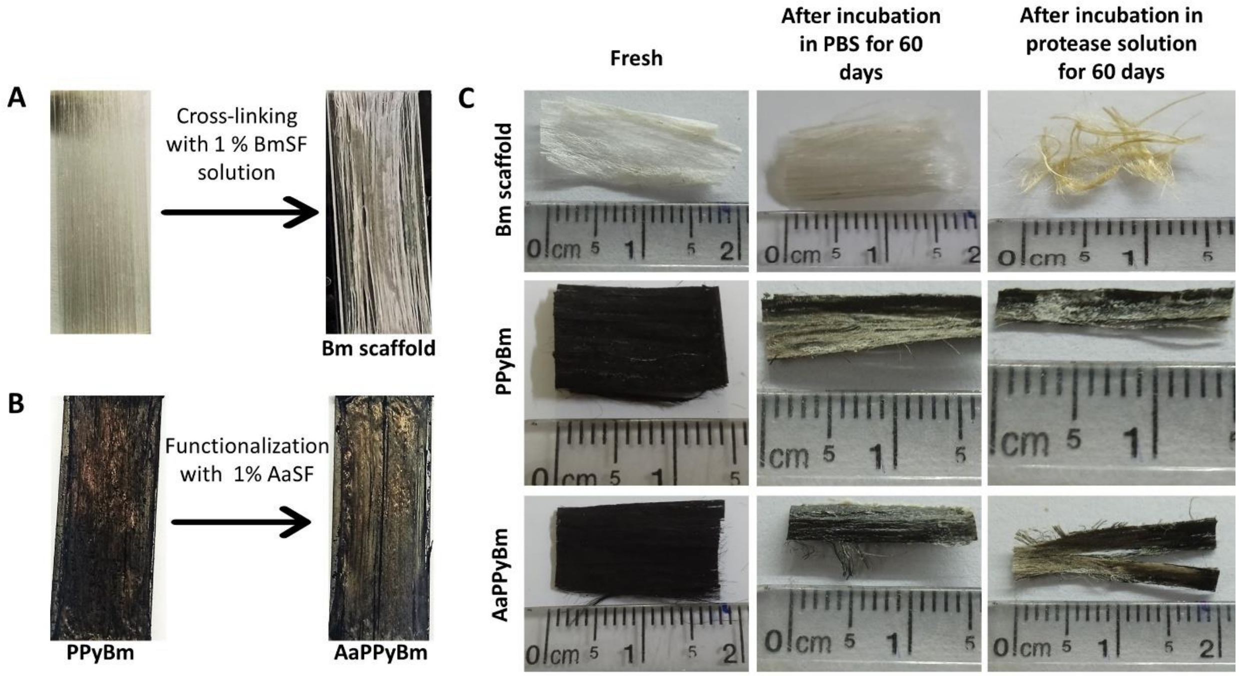


**Figure S9**: Photographs of aligned (A) degummed Bm fibers after cross-linking with regenerated BmSF and (B) PPy:Silk scaffold after functionalization with regenerated AaSF. C. Representative images of fresh and degraded scaffolds (after incubation in PBS and protease solution for 60 days) as indicated.

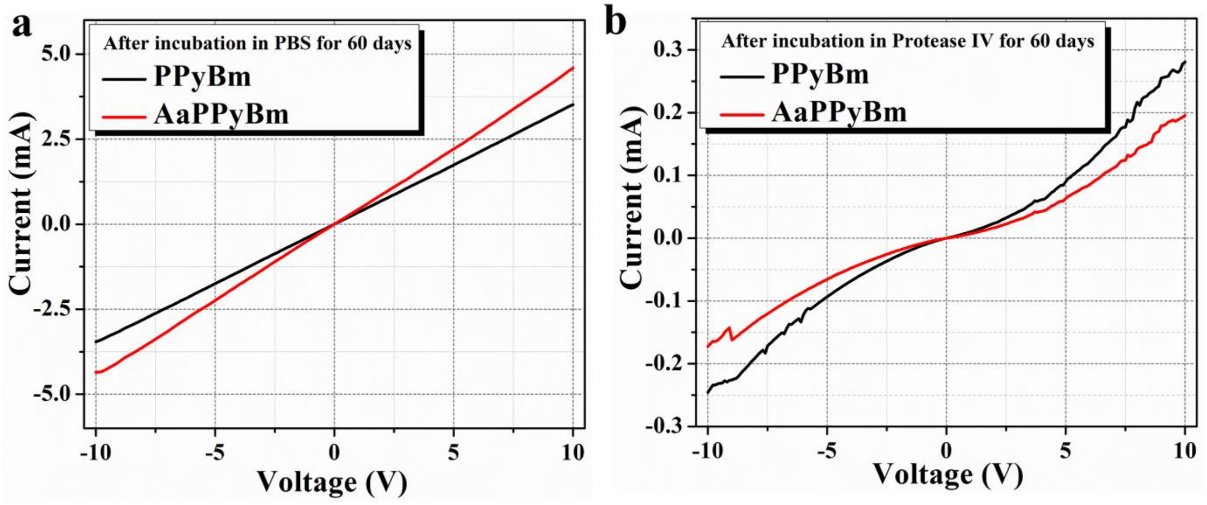


**Figure S10**: Current-voltage (I-V) characteristics of various PPy:Silk based scaffolds recorded after 60 day incubation in (A) PBS and (B) protease IV.

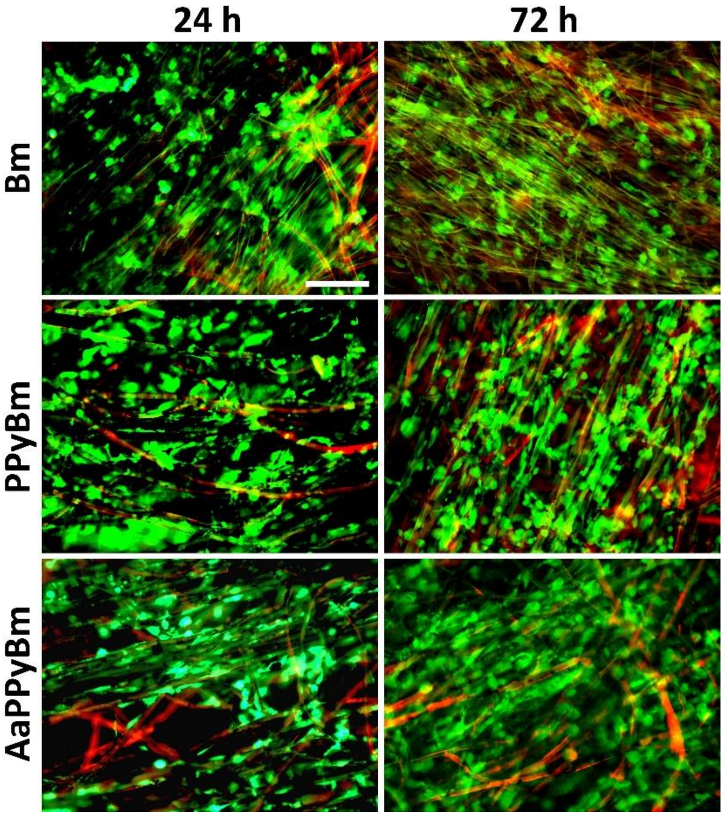


**Figure S11**: Preliminary in-vitro biocompatibility studies of various silk-based scaffolds as indicated. Representative live-dead staining images showing viability of MG63 cells on different scaffolds after 24 and 72 h. Calcein-AM stained live cells appear as green, while ethidium homodimer (EthD-1) stained dead cells can be seen in red. Red autofluorescence from scaffold depicts their aligned morphology; Scale bar = 200 μm.

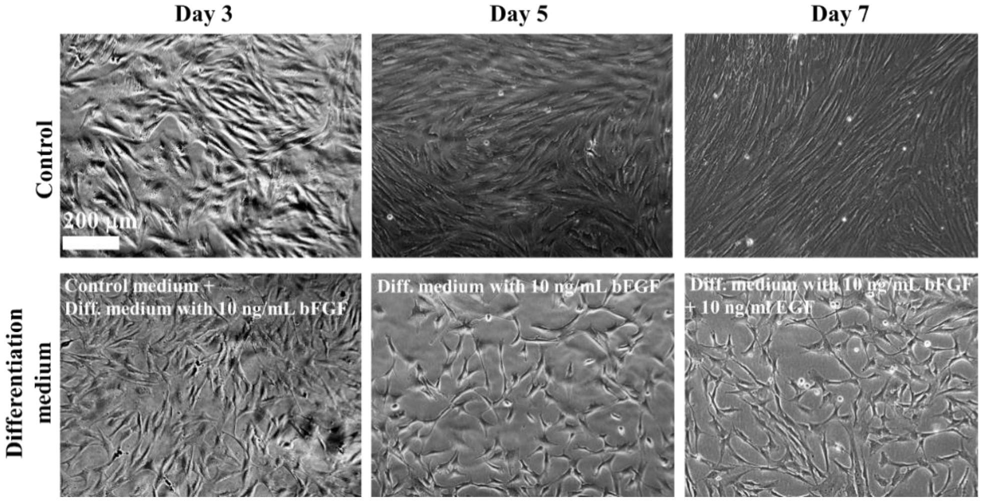


**Figure S12**: Representative phase-contrast micrographs showing neural differentiation of primary adipose derived mesenchymal stem cells (pADSCs). Morphological changes of cells at different steps of differentiation process when compared with morphology of cells cultured in control medium (DMEM + 10% FBS+ 1% penicillin-streptomycin). Neurite like extensions started to appear after Day 5 after treating with complete differentiation media (DMEM + 10% FBS+ 1% penicillin-streptomycin+ 10 ng/mL bFGF+ 10 ng/mL EGF).

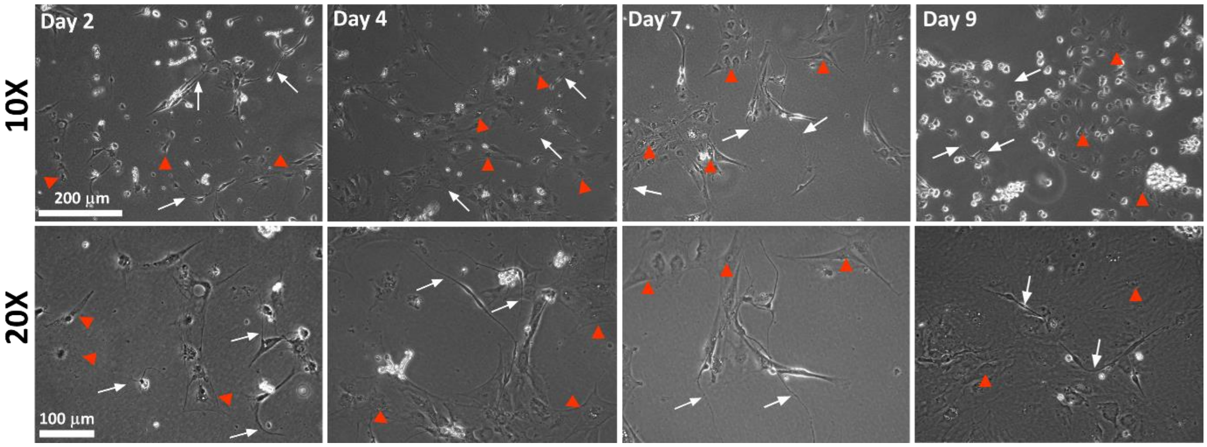


**Figure S13**: Representative phase-contrast micrographs of dissociated pDRG culture after isolation showing different cell morphologies at different time-period. Though, the culture consists of mixed cell populations, neurons with neurite projections (white arrow) are clearly visible. Other than sensory neurons, pDRG also contains other cell types such as satellite glial cells, Schwann cells, fibroblasts etc.,[17, 20, 21] which are also indicated with red arrowhead. The density of other cell types decreases with culture time in neuron specific media.

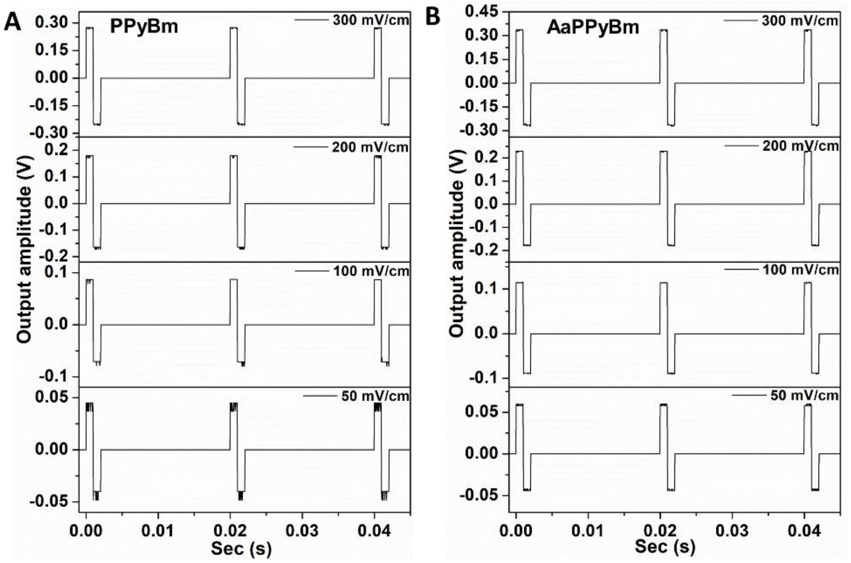


**Figure S14**: Representative pulsed waveforms recorded using an oscilloscope during electrical stimulation (ES) of DRG neurons through the non-functionalized and functionalized PPy:Silk scaffolds at different amplitudes.

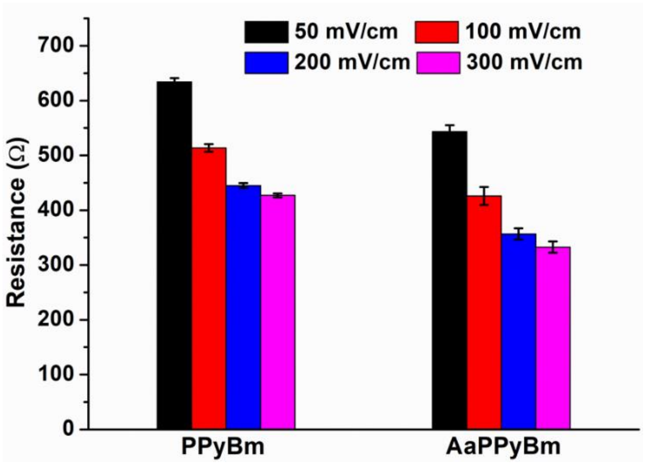


**Figure S15**: Surface resistance exhibited by the PPy:Silk based scaffolds at different bias voltages as labelled.
